# The neural representation of visually evoked emotion is high-dimensional, categorical, and distributed across transmodal brain regions

**DOI:** 10.1101/872192

**Authors:** Tomoyasu Horikawa, Alan S. Cowen, Dacher Keltner, Yukiyasu Kamitani

**Affiliations:** Department of Neuroinformatics, ATR Computational Neuroscience Laboratories, Hikaridai, Seika, Soraku, Kyoto, 619-0288, Japan; Department of Psychology, University of California, Berkeley, CA, 94720-1500, USA; Google Research, Mountain View, CA, 94043, USA; Graduate School of Informatics, Kyoto University, Yoshida-honmachi, Sakyo-ku, Kyoto, 606-8501, Japan; Lead contact

**Keywords:** emotion categories, affective dimensions, decoding, encoding, functional magnetic resonance imaging

## Abstract

Central to our subjective lives is the experience of different emotions. Recent behavioral work mapping emotional responses to 2185 videos found that people experience upwards of 27 distinct emotions occupying a high-dimensional space, and that emotion categories, more so than affective dimensions (e.g., valence), organize self-reports of subjective experience. Here, we sought to identify the neural substrates of this high-dimensional space of emotional experience using fMRI responses to all 2185 videos. Our analyses demonstrated that (1) dozens of video-evoked emotions were accurately predicted from fMRI patterns in multiple brain regions with different regional configurations for individual emotions, (2) emotion categories better predicted cortical and subcortical responses than affective dimensions, outperforming visual and semantic covariates in transmodal regions, and (3) emotion-related fMRI responses had a cluster-like organization efficiently characterized by distinct categories. These results support an emerging theory of the high-dimensional emotion space, illuminating its neural foundations distributed across transmodal regions.

## Introduction

Emotions are mental states generated by the brain, constituting an evaluative aspect of our diverse subjective experiences. Traditionally, subjective emotional experiences have been described in theoretical and empirical studies using a limited set of emotion categories (e.g., happiness, anger, fear, sadness, disgust, and surprise) as proposed in early versions of basic emotion theory (Ekman et al., 1969; Plutchik, 1980) and by broad affective dimensions (e.g., valence and arousal) that underpin the circumplex model of affect and the more recent core affect theory (Posner et al., 2005; Russell, 1980, 2003). Grounded in these theoretical claims, affective neuroscientists have investigated neural signatures underlying those representative emotion categories and affective dimensions (Barrett, 2017; Celeghin et al., 2017; Giordano et al., 2018; Hamaan, 2012; Kragel and LaBar, 2015; Lindquist and Barrett, 2012; Saarimäki et al., 2016; Satpute and Lindquist, 2019; Wager et al., 2015), finding that distributed brain regions and functional networks are crucial for representing individual emotions. Importantly, though, studies within this tradition have limited their focus to a small set of emotions. This narrow focus does not capture the increasingly complex array of states associated with distinct experiences (Cowen and Keltner, 2017, 2019; Cowen et al., 2018, 2019a, 2020). As a result, much of the variability in the brain’s representation of emotional experience likely has yet to be explained (Cowen et al., 2019a; Lindquist and Barrett, 2012).

To elucidate the structure of diverse emotional experiences, Cowen and Keltner (2017, 2019; Cowen et al., 2018, 2019a, 2020) offered a new conceptual and methodological approach to capture the more complex “high-dimensional emotion space” that characterizes emotional experiences in response to a diverse array of stimuli in large-scale human behavioral experiments. In their study, Cowen and Keltner (2017) applied statistical methods to analyze reported emotional states elicited by 2185 videos each annotated by multiple raters using self-report scales of 34 emotion categories (e.g., joy, amusement, and horror) and 14 affective dimensions (e.g., valence and arousal) derived from basic, constructivist, and dimensional theories of emotion. They identified 27 distinct kinds of emotional experience reliably associated with distinct videos. Furthermore, they found that the affective dimensions explained only a fraction of the variance in self-reports of subjective experience when compared to emotion categories, a finding that has now been replicated in studies of facial expression, speech prosody, nonverbal vocalization, and music (Cowen and Keltner, 2019; Cowen et al., 2018, 2019a, 2020). This work lays a foundation for examining the neural bases of emotional experience in terms of a rich variety of states, as well as whether — at the level of neural response — categories or dimensions organize the experience of emotion, and how a diverse array of states cluster in their neural representations.

The methodological criteria established in this study of emotion — the use of broad ranging stimuli sufficient to compare theoretical models of emotion — can be readily extended to the study of the neural basis of emotion. In addition, it is important for studies of the neural basis of emotion to consider activity across the whole brain, given that diverse psychological processes and distributed brain regions and functional networks are known to contribute to the representation of various emotional states (Barrett, 2017; Celeghin et al., 2017; Hamaan, 2012; Lindquist and Barrett, 2012; Satpute and Lindquist, 2019). Although some recent studies have examined a richer variety of emotions than typically tested before (more than twenty), their investigations have been focused on specific brain regions (Kragel et al., 2019; Skerry and Saxe, 2015) and have relied on relatively small numbers of stimuli or conditions (tens or hundreds, but not thousands) (Koide-Majima, et al., 2018; Saarimäki et al., 2018). Moreover, studies have not adequately differentiated representations of emotion from representations of sensory and semantic features that are known to be encoded throughout the cortex (Huth et al., 2012) and may contaminate neural responses to emotional stimuli (Kragel et al., 2019; Skerry and Saxe, 2015). Given these considerations, a comprehensive analysis of whole-brain responses to diverse emotional stimuli is still needed.

We sought to understand the neural underpinnings of the high-dimensional space of diverse emotional experiences associated with a wide range of emotional stimuli using the experimental resources generated in the study by Cowen and Keltner (2017). Specifically, we measured whole-brain functional magnetic resonance imaging (fMRI) signals while five subjects watched 2181 emotionally evocative short videos. We then analyzed the measured fMRI signals using data-driven approaches, including neural decoding, voxel-wise encoding, and unsupervised modeling methodologies (Figure 1A) to investigate neural representations of emotional experiences characterized by 34 emotion categories and 14 affective dimensions with which the videos were previously annotated by a wide range of raters (Figure 1B).

**Figure 1.**
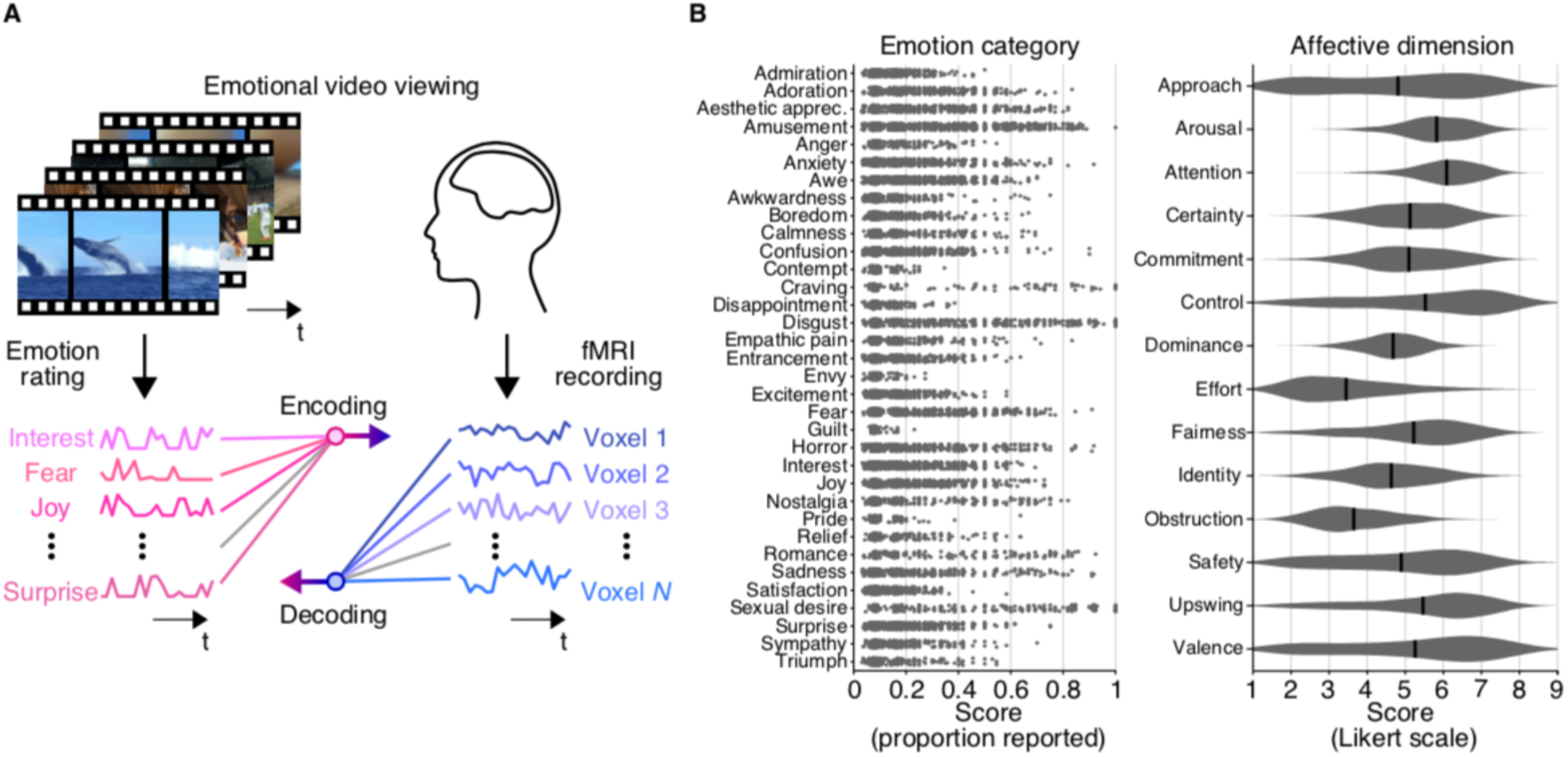
Decoding and encoding emotional responses in fMRI signals evoked by videos. (A) Schematic of the experiment and analyses. fMRI signals were recorded while subjects viewed emotional videos (with no sound). Decoding/encoding models were trained to predict scores/responses of individual emotions/voxels for presented videos from patterns of responses/scores, respectively. (B) Score distributions of emotion categories and affective dimensions. Score distributions from a total of 2181 unique videos are shown for 34 emotion categories and 14 affective dimensions (see Transparent Methods: “Video stimulus labeling”; Cowen and Keltner, 2017 for more details). For emotion categories, each dot indicates a proportion that the category was evaluated by raters with non-zero scores for each video. For visualization purposes, only dots corresponding to videos with non-zero scores were shown with jittering. For affective dimensions, black bars indicate mean values across all videos. In the main analyses, mean scores averaged across multiple raters were used for each video.

Using multiple analytical approaches, we investigated different facets of the neural representation of the emotional experiences associated with the videos. Neural decoding was used to examine whether and where individual features (e.g., ratings of an emotion) are represented in the brain, and provides a powerful way to identify mental states of subjects from brain activity patterns (Horikawa and Kamitani, 2017; Kragel et al., 2016; Sitaram et al., 2011). Voxel-wise encoding modeling made it possible to evaluate how specific sets of features (e.g., emotion category scores) modulate activity of individual voxels, and to characterize representational properties of individual voxels by comparing encoding accuracies from different feature sets (de Heer et al., 2017; Güçlü and van Gerven, 2015; Kell et al., 2018; Lescroart and Gallant, 2019). Unsupervised modeling methods, including dimensionality reduction and clustering analyses, were used to characterize the distributional structure of brain representations of emotion in an exploratory manner (Kriegeskorte et al., 2008).

Here, we first show that dozens of distinct emotions associated with videos were reliably predicted from activity patterns in multiple brain regions, which enabled accurate identification of emotional states. While the brain regions representing individual emotions overlapped, configurations of regions proved to be unique to each emotion and consistent across subjects. We show that voxel activity in most brain regions was better predicted by voxel-wise encoding models trained on ratings of emotion categories than on ratings of affective dimensions. Comparisons with other models trained on visual or semantic features revealed that the emotion category model uniquely explained response modulations of voxels in transmodal regions, indicating that these regions encode either emotional experience or visual affordances for specific emotional experiences (e.g., the inference that a landscape is likely to inspire awe). Finally, unsupervised modeling of emotion-related neural responses showed that distributions of brain activity patterns associated with diverse emotional experiences (or affordances) were organized with clusters corresponding to specific emotion categories, which were bridged by continuous gradients. These results provide support for the theory that emotion categories occupy a high-dimensional space (Cowen and Keltner, 2017) and shed new light on the neural underpinnings of emotion.

## Results

To understand neural representations of diverse emotional experiences associated with rich emotional content, we recorded whole-brain fMRI signals from five subjects while they were freely viewing emotionally evocative short videos (Figure 1A). The stimuli consisted of a total of 2181 unique videos annotated with a total of 48 emotion ratings (Figure 1B; 34 emotion categories and 14 affective dimensions), each by multiple human raters, which were collected in a previous study (Cowen and Keltner, 2017; see Transparent Methods: “Video stimulus labeling”). The fMRI volumes measured over the course of each video presentation were shifted by 4 s and averaged to estimate fMRI responses to individual video stimuli. We performed decoding and encoding analyses between the fMRI responses to each video and its emotion ratings by using regularized linear regression in a cross-validated manner (6-fold cross-validation; see Transparent Methods: “Regularized linear regression analysis”). We also performed unsupervised modeling of the fMRI responses to examine the distribution of emotion-related brain activity patterns and whether their organization was better described by emotion categories or by broad affective dimensions (see Transparent Methods: “Dimensionality reduction analysis” and “Clustering analysis”).

### Neural decoding analysis to predict emotion scores

We first used a neural decoding analysis to examine whether the emotions associated with presented videos could be predicted from brain activity patterns within specific regions and across the brain. We constructed a decoding model (decoder) for each emotion (34 emotion categories and 14 affective dimensions) to predict emotional experience ratings from the brain using multi voxel activity patterns as input. The decoders of individual emotions were separately trained using activity patterns in each of multiple brain regions that include 360 cortical regions defined by a parcellation provided from the Human Connectome Project (Glasser, et al., 2016; 180 cortical areas per hemisphere; HCP360) and 10 subcortical regions (e.g., amygdala and cerebellum). This procedure thus yielded a total of 370 region-wise decoders from brain activity per emotion. The performance for each emotion and brain region was evaluated by computing a Pearson correlation coefficient between predicted and true emotion scores across the 2181 unique videos.

The decoding accuracies for the categories and dimensions obtained from multiple brain regions are shown in Figures 2A and B, respectively (see Figure 2C for color/shape schema of individual brain regions; also see Figures S1A and B for results of representative cortical regions and subcortical regions). Although scores of three emotion categories (envy, contempt, and guilt) were not reliably predicted from any brain regions consistently across five subjects, scores of the remaining emotions were predicted from brain activity patterns in at least one brain region with significant accuracy (*r* > 0.095, permutation test, *p* < 0.01, Bonferroni correction by the number of brain regions) — typically, from many different regions as the color of individual dots (or brain regions) indicated. The results established that ratings of the 27 emotion categories that were reported to be reliably elicited by video stimuli (Cowen and Keltner, 2017) were highly decodable from multiple brain regions. The poor accuracies for a few categories of emotion may be attributable in part to relatively lower proportions of videos evoking these categories (Figure 1B).

**Figure 2.**
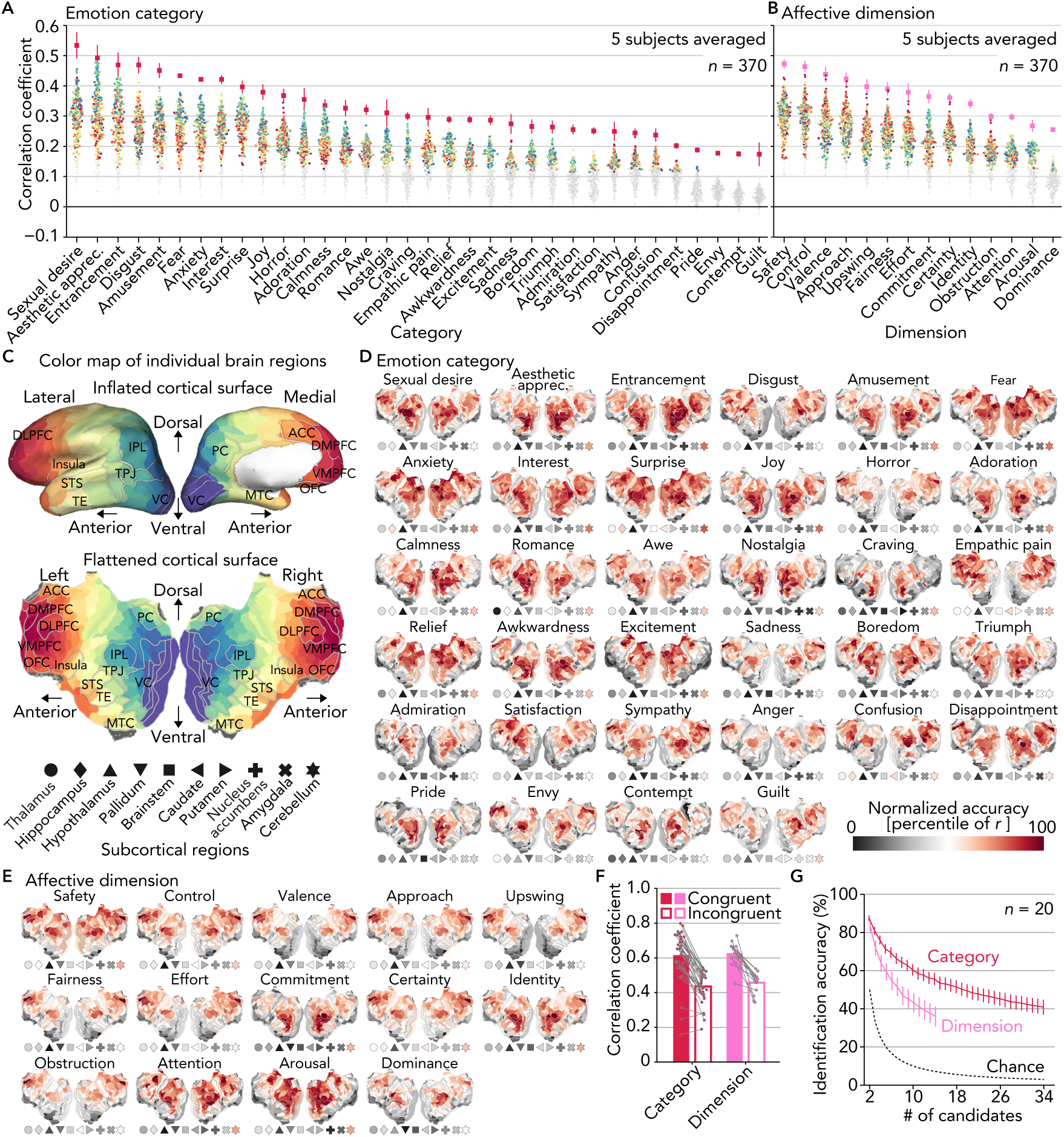
Neural decoding analysis to predict emotion scores. (A) Decoding accuracy for individual categories. Dots indicate accuracies obtained from individual cortical regions of the HCP parcellation and subcortical regions (*n* = 370, five subjects averaged; see Figures S1A and B for results of representative cortical regions and subcortical regions). Red/pink squares indicate accuracies by ensemble decoders that aggregate predictions from multiple brain regions (error bars, 95% confidence intervals [C.I.] across subjects). Brain regions with significant accuracy from all five subjects are colored (*r* > 0.095, permutation test, *p* < 0.01, Bonferroni correction by the number of brain regions). Color and shape of each dot indicate locations of individual brain regions as (C). (B) Decoding accuracy for individual dimensions. Conventions are the same as (A). (C) Color map of individual brain regions. Several brain regions are outlined and labeled (abbreviations: VC, visual cortex; TPJ, temporo-parietal junction; IPL, inferior parietal lobule; PC, precuneus; STS, superior temporal sulcus; TE, temporal area; MTC, medial temporal cortex; DLPFC/DMPFC/VMPFC, dorsolateral/dorsomedial/ventromedial prefrontal cortex; ACC, anterior cingulate cortex; and OFC, orbitofrontal cortex). (D) Cortical surface maps of decoding accuracies for individual categories (five subjects averaged). Conventions of the marker points for the subcortical regions are the same as (C). (E) Cortical surface maps of decoding accuracies for individual dimensions (five subjects averaged). Conventions are the same as (D). (F) Correlations of region-wise decoding accuracy patterns between paired subjects. In congruent conditions, each dot indicates a mean correlation between the same emotion pairs averaged across all pairs of subjects. In incongruent conditions, each dot indicates a mean correlation between different emotion pairs averaged across all pairs of subjects and emotions. (G) Emotion identification accuracy based on inter-individual similarities of region-wise decoding accuracy patterns (error bars, 95% C.I. across pairs of subjects [*n* = 20]).

To integrate information represented in multiple brain regions, we constructed an ensemble decoder for each emotion by averaging predictions from multiple region-wise decoders selected based on decoding accuracy via a nested cross-validation procedure (red and pink squares in Figure 2A and B, respectively; see Transparent Methods: “Construction of ensemble decoders”). Overall, the ensemble decoders tended to show higher accuracy than the best region-wise decoders (143/170 = 84.1% for categories, 52/70 = 74.3% for dimensions, pooled across emotions and subjects). These results indicate that information about individual emotions is represented in multiple brain regions in a complementary manner, supporting the notion of distributed, network-based representations of emotion (Barrett, 2017; Celeghin et al., 2017; Hamaan, 2012; Lindquist and Barrett, 2012; Saarimäki et al., 2016; Satpute and Lindquist, 2019).

### Region-wise decoding accuracy maps

To investigate which brain regions represented individual emotions, we visualized decoding accuracy maps by projecting accuracies in the 360 cortical regions onto flattened cortical surfaces alongside markers representing accuracies in the ten subcortical regions (Figures 2D and E; five subjects averaged). In general, brain regions showing high accuracy for specific emotions were not focal but rather distributed across multiple regions including visual, parietal, and frontal regions. For instance, scores of several basic emotions like sadness and anger were well predicted from distant regions around parietal (IPL) and frontal (DMPFC) regions (different hubs of default mode network). These results further support the notion that distributed brain regions and functional networks are recruited to represent individual emotions.

Together with the color differences of accuracy distributions across emotions in Figures 2A and B, visual inspections of the accuracy maps revealed different configurations of highly informative brain regions for different emotions. Interestingly, even subjectively related emotions (e.g., fear and horror, confusion and awkwardness) were often represented in markedly different configurations of brain regions, supporting a nuanced taxonomy of emotion.

### Inter-individual consistency in brain regions representing distinct emotions

To examine the extent to which the configurations of brain regions informative for decoding were unique to each emotion and consistent across subjects, we next tested whether emotions could be identified across subjects based on the region-wise decoding accuracy patterns of individual emotions estimated for individual subjects. A region-wise accuracy pattern for one emotion from one subject was used to identify the same emotion among a variable number of candidates (a pattern for the same emotion and randomly selected patterns for other emotions) from another subject using Pearson correlation coefficients (see Transparent Methods: “Emotion identification analysis”). The analysis was performed for all pairs among five subjects (*n* = 20) separately using the accuracies for the categories and dimensions. For most of the categories and dimensions, correlations between the same emotion pairs (congruent; mean correlations, 0.615 for categories, 0.628 for dimensions) were larger than correlations between different emotion pairs (incongruent; mean correlations, 0.435 for categories, 0.458 for dimensions) (Figure 2E). Furthermore, mean identification accuracy averaged across all pairs of subjects far exceeded chance levels across all candidate set sizes for both of the categories and dimensions (Figure 2F; e.g., 5-way identification accuracy, 71.4% for categories, 59.3% for dimensions; chance level, 20%). These results suggest that distinct emotions associated with the video stimuli were differentially represented across the brain in a consistent manner across individuals.

### Identification of videos via decoded emotion scores

Given that a complex emotional state can be represented by a blend or combination of multiple emotions (e.g., anger with sadness), we next tested a decoding procedure that took into account multiple emotions simultaneously. To do so, we examined whether a specific video stimulus could be predicted, or identified, from the concatenated predictions of emotion categories or those of affective dimensions. The identification analysis was performed in a pairwise manner, in which a vector of predicted scores was compared with two candidate vectors (one from a true video and the other from a false video randomly selected from the other 2180 videos) to test whether a correlation for the true video was higher than that for the false video (see Transparent Methods “Video identification analysis”). For each video, the analysis was performed with all combinations of false candidates (*n* = 2180), and percentages of correct answers were used as an identification accuracy for one sample.

Identification accuracies obtained from predictions by the region-wise decoders are shown in Figure 3A (five subjects averaged, see Figure S1C for individual subjects). The results showed better identification accuracy via the categories than via the dimensions with most region-wise decoders. The superior accuracy of the categories over the dimensions was also observed with the ensemble decoders, which showed greater than 10% differences in mean accuracy in all subjects (Figure 3B; 81.9% via categories, 68.7% via dimensions, five subjects averaged; see Figure 3C for accuracies of individual videos). This tendency was preserved even when the number of emotions used for identification was matched between the category and dimension models by randomly selecting a subset of 14 categories from the original 34 categories (Figure S1D; 75.4%, five subjects averaged). These results indicate that emotion categories capture far more variance, and presumably details, in the neural representation of emotionally evocative videos than broad affective features such as valence and arousal.

**Figure 3.**
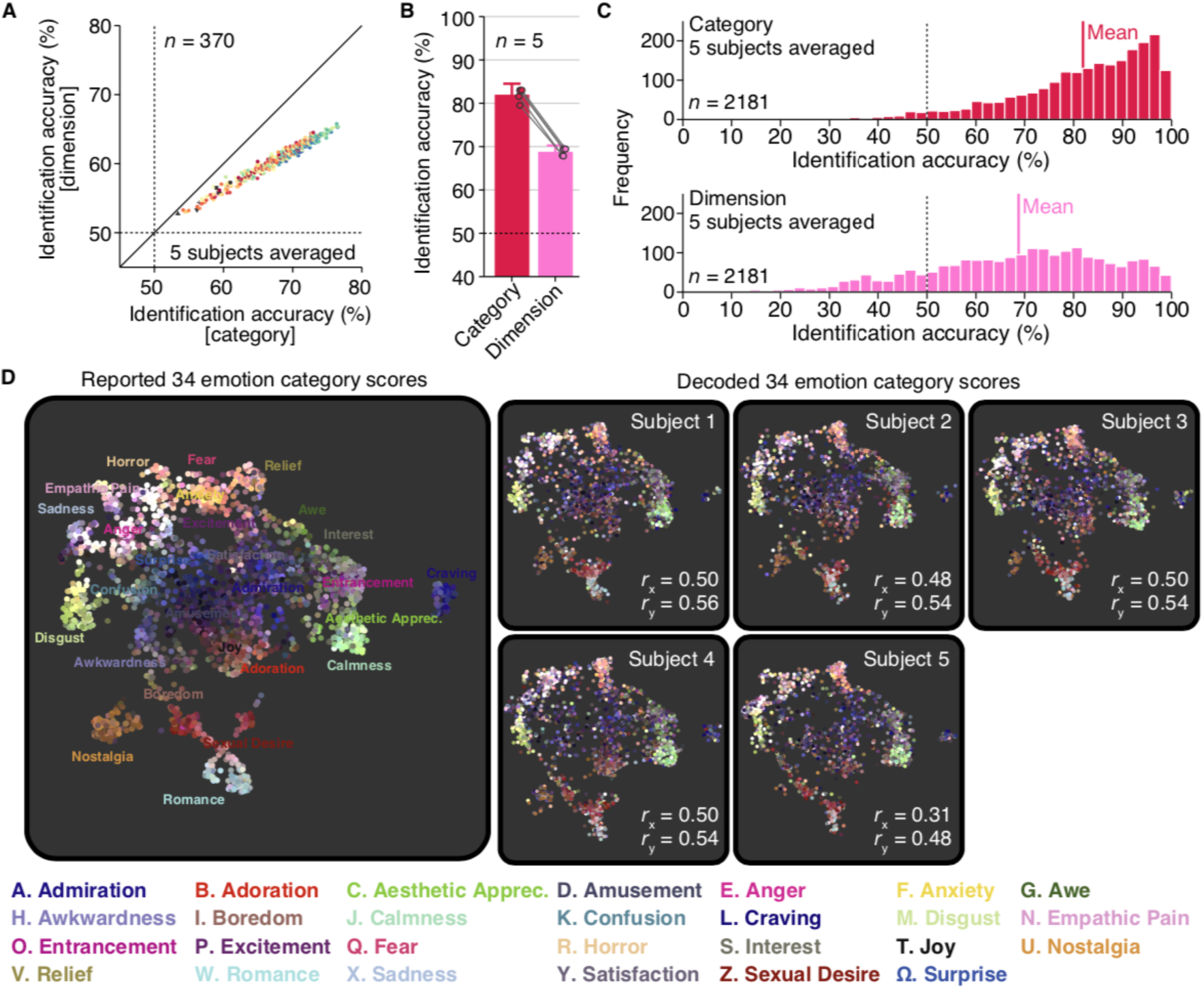
Identification of videos via decoded emotion scores. (A) Mean video identification accuracies from region-wise decoders. Dots indicate accuracies from individual brain regions (five subjects averaged; dashed lines, chance level, 50%; see Figure S1C for individual subjects). Color and shape of each dot indicate locations of individual brain regions as Figure 2C. (B) Mean video identification accuracies from ensemble decoders. Dots indicate accuracies of individual subjects (error bars, 95% C.I. across subjects; dashed line, chance level, 50%). (C) Histogram of video identification accuracies for individual videos (ensemble decoder; five subjects averaged; dashed line, chance level, 50%). (D) Two-dimensional maps of emotional experiences constructed from reported and decoded scores. Each dot corresponds to each video, and was colored according to its original reported scores using a weighted interpolation of unique colors assigned to 27 distinct categories (see Cowen and Keltner, 2017 for the color schema).

### Visualization of decoded emotional experiences (or affordances)

To explore the extent to which richly varying emotional states were accurately decoded, we used a nonlinear dimensionality reduction method to visualize the distribution of emotional experiences evoked by the 2181 videos, originally represented in terms of 34 emotion categories, in a two-dimensional space (Cowen and Keltner, 2017), and tested whether the map could be reconstructed from the decoded scores. For this purpose, we used a novel dimensionality reduction algorithm, called Uniform Manifold Approximation and Projection (UMAP; McInnes et al., 2018), which enables the projection of new data points after constructing a mapping function from an independent training dataset. We first constructed a mapping function that projects 34-dimensional emotion category scores into a two-dimensional space using the original emotion category scores (Figure 3D left), and then the function was used to project decoded scores from ensemble decoders of individual subjects onto the map (Figure 3D right).

We first confirmed that the UMAP algorithm applied to the 34 emotion category scores replicated a similar two-dimensional map produced in the previous study (Cowen and Keltner, 2017), in which distinct categories were found to be organized along gradients between clusters associated with each category. Upon inspection of the maps reconstructed from decoded scores, it is apparent that distributions of colors (scores) of the original map were precisely replicated for all subjects. This was confirmed quantitatively by calculating correlations of horizontal (x) and vertical (y) data positions in the two-dimensional space between true and reconstructed maps (*r* = 0.46 for horizontal axis, *r* = 0.53 for vertical axis, five subjects averaged). The results demonstrate how precisely different emotional states were decoded from brain activity and provide a useful means to visualize decoded emotional states.

### Construction of voxel-wise encoding models

We next used voxel-wise encoding models to characterize how varying features associated with the presented videos modulate activities in each voxel. We constructed an encoding model for each voxel to predict activities induced by the emotionally evocative videos using a set of features (e.g., scores for 34 emotion categories), and model performance was evaluated by computing a Pearson correlation coefficient between measured and predicted activities (see Transparent Methods: “Regularized linear regression analysis”).

### Encoding models predicting brain activity from emotional scores

We first tested how well encoding models constructed from the 34 emotion categories (e.g., joy, anger) and the 14 affective dimensions (e.g., valence, arousal) predicted brain activities induced by the presented videos (Figure 4A; Subject 1; see Figure S2A for the other subjects). We found that both the category and dimension models accurately predicted activity in many regions, indicating that a broad array of brain regions are involved in encoding information relevant to emotion, as represented by categories and affective dimensions.

**Figure 4.**
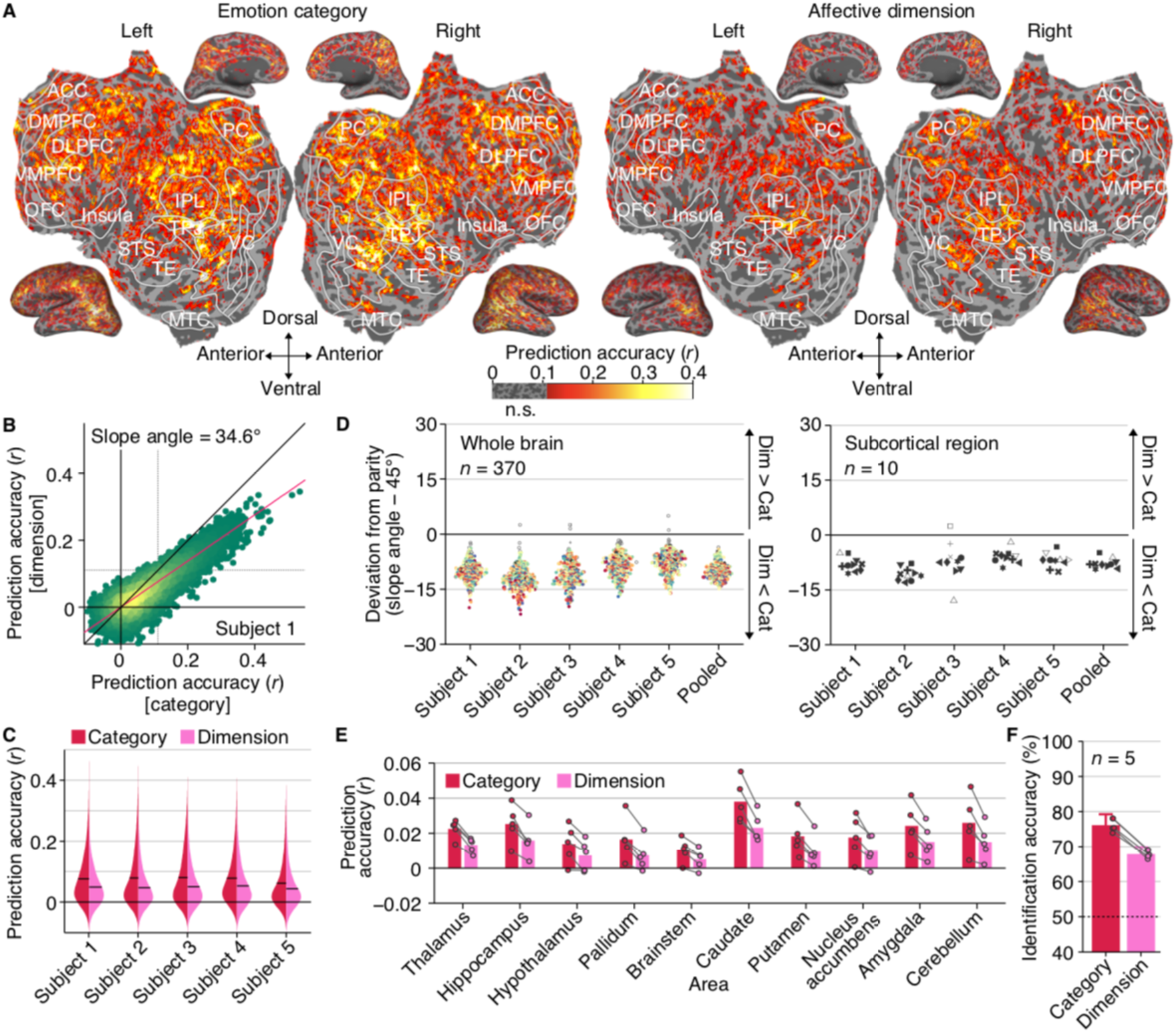
Encoding models predicting brain activity from emotional scores. (A) Prediction accuracies of emotion encoding models (Subject 1, see Figure S2A for the other subjects). (B) Prediction accuracies of individual voxels (red line, best linear fit; dotted lines, *r* = 0.111, permutation test, *p* < 0.01, Bonferroni correction by the number of voxels; see Figure S2B for the other subjects). (C) Distributions of prediction accuracies of all voxels (black bars, mean of all voxels). (D) Comparisons of prediction accuracies for individual brain regions. Each dot indicates a deviation of the estimated slope angle from the parity for each brain region (baseline, 45 degree). Brain regions with significant deviation are colored/filled (two-tailed t-test, *p* < 0.01 by jackknife method, Bonferroni correction by the number of ROIs, *n* = 370). Color and shape of each dot indicate locations of individual brain regions as Figure 2C. (E) Mean prediction accuracies in individual subcortical regions. Dots indicate accuracies of individual subjects. (F) Mean video identification accuracy via predicted brain activities. Dots indicate accuracies of individual subjects (error bars, 95% C.I. across subjects; dashed line, chance level, 50%).

In comparing the model performances, we found that on average the category model outperformed the dimension model by a considerable margin (Figure 4B; Subject 1; slope angle, 34.6 degree; two-tailed t-test, *p* < 0.01 by jackknife method; see Figure S2B for the other subjects). In comparison with the dimension model, the category model predicted voxel responses significantly better in all subjects (Figure 4C, paired t-test across all voxels; *p* < 0.01 for all subjects), and also predicted 73.0% more voxels significantly predicted voxels (*r* > 0.111, permutation test, *p* < 0.01, Bonferroni correction by the number of voxels; five subjects averaged). In addition, when we constructed a joint encoding model by concatenating scores of 34 categories and 14 dimensions, the increase in the number of significantly predicted voxels was marginal when compared to the category model (1.8%, five subjects averaged), but substantial when compared to the dimension model (75.9%, five subjects averaged). Together, these findings suggest that the brain representation of affective dimensions is subsumed by, and can be inferred from, the brain representation of emotion categories.

We next explored whether there were particular brain regions in which the dimension model outperformed the category model. We divided the whole cortex into 360 regions using the HCP360 parcellation (Glasser et al., 2016), and then separately compared the performances in each of the 360 cortical and 10 subcortical regions using a slope angle of best linear fit of the accuracies from the two models (see Transparent Methods: “Slope estimates for performance comparisons”). The distributions of the slope angle suggest that activity in nearly every region was better predicted by the category model (Figure 4D left; 368/370 for category, 0/370 for dimension; two-tailed t-test, *p* < 0.01 by jackknife method, Bonferroni correction by the number of ROIs, *n* = 370; five subjects pooled). Notably, this tendency was even observed in ten subcortical regions, including the amygdala (see Transparent Methods: “Regions of interest (ROI)”), which are often thought to prioritize simpler appraisals of valence and arousal (Figures 4D right; 9/10 for category, 0/10 for dimension; two-tailed t-test, *p* < 0.01 by jackknife method, Bonferroni correction by the number of ROIs, *n* = 370; five subjects pooled; see Figure 4E for mean encoding accuracy for the subcortical regions).

To further compare the performance of the category and dimension models, a video identification analysis was performed via predicted brain activities obtained from each of the two models (cf., Figure 3B; see Transparent Methods: “Video identification analysis”). We confirmed that identification accuracies from the category model were substantially better than the dimension model for all subjects (Figure 4F; 76.1% for category, 67.9% for dimension, five subjects averaged).

In sum, these encoding analyses demonstrate that emotion categories substantially outperform affective dimensions in capturing voxel activity induced by emotional videos (see Figure S3 for control analyses of potential confounds). Taken together, these results build on previous behavioral findings (Cowen and Keltner, 2017), which showed that self-reported emotional experience was more precisely characterized by a rich array of emotion categories than by a range of broad affective dimensions proposed in constructivist and componential theories to be the underlying components of emotional experience (Barrett, 2017; Posner et al., 2005; Smith and Ellsworth, 1985). In particular, using a wide range of analytic approaches, we repeatedly found that emotion categories capture nuanced, high-dimensional neural representations that cannot be accounted for by a lower-dimensional set of broad affective features.

### Disentangling emotional, visual object, and semantic feature encoding

Given that our video stimuli consisted of scenes and events that also included many visual objects and semantic features, we next sought to disentangle brain representations of emotion from those correlated features. For this analysis, we constructed two additional encoding models from outputs of a pre-trained deep neural network for object recognition (“visual object model”; Simonyan and Zisserman, 2014) and from semantic features (or concepts; e.g., cats, indoor, and sports) annotated by human raters via crowdsourcing (“semantic model”; see Transparent Methods: “Video stimulus labeling” for these features). We then evaluated model performances for each model and compared them to those of the “emotion model” constructed from the emotion category scores.

As found in previous studies that used similar visual (Güçlü and van Gerven, 2015) and semantic (Huth et al., 2012) encoding models, our visual object model and semantic model were also effective in predicting voxel activity elicited by video stimuli even outside of the visual cortex (Figure 5A; Subject 1; see Figure S4A for the other subjects). While the accuracies from these two models positively correlated with those from the emotion model (inset in Figure 5A), the emotion model consistently outperformed the other two models in certain brain regions (Figure 5B), including IPL, TE, and several frontal regions (DLPFC, DMPFC, VMPFC, and OFC), which are known to represent multimodal emotion information (Chikazoe et al., 2014; Skerry and Saxe, 2014) and abstract semantic features (Huth et al., 2016; Skerry and Saxe, 2015), and in multiple functional networks (Figure 5C; Yeo et al., 2011), including the salience network (8), subparts of the limbic network (10), executive control network (13), and default mode network (16 and 17), which is also consistent with previous reports (Barrett, 2017; Barrett and Satpute, 2013; Lindquist and Barrett, 2012; Satpute and Lindquist, 2019). Furthermore, while accuracies were not as high, activity in the subcortical regions also tended to be better predicted by the emotion models than by the other two models (Figure 5D). This analysis demonstrates that brain regions preferentially encoding emotion are broadly distributed in a similar manner across subjects (Figure 5E; see Figure S4D for evaluation based on slope estimates, cf., Figure 4C).

**Figure 5.**
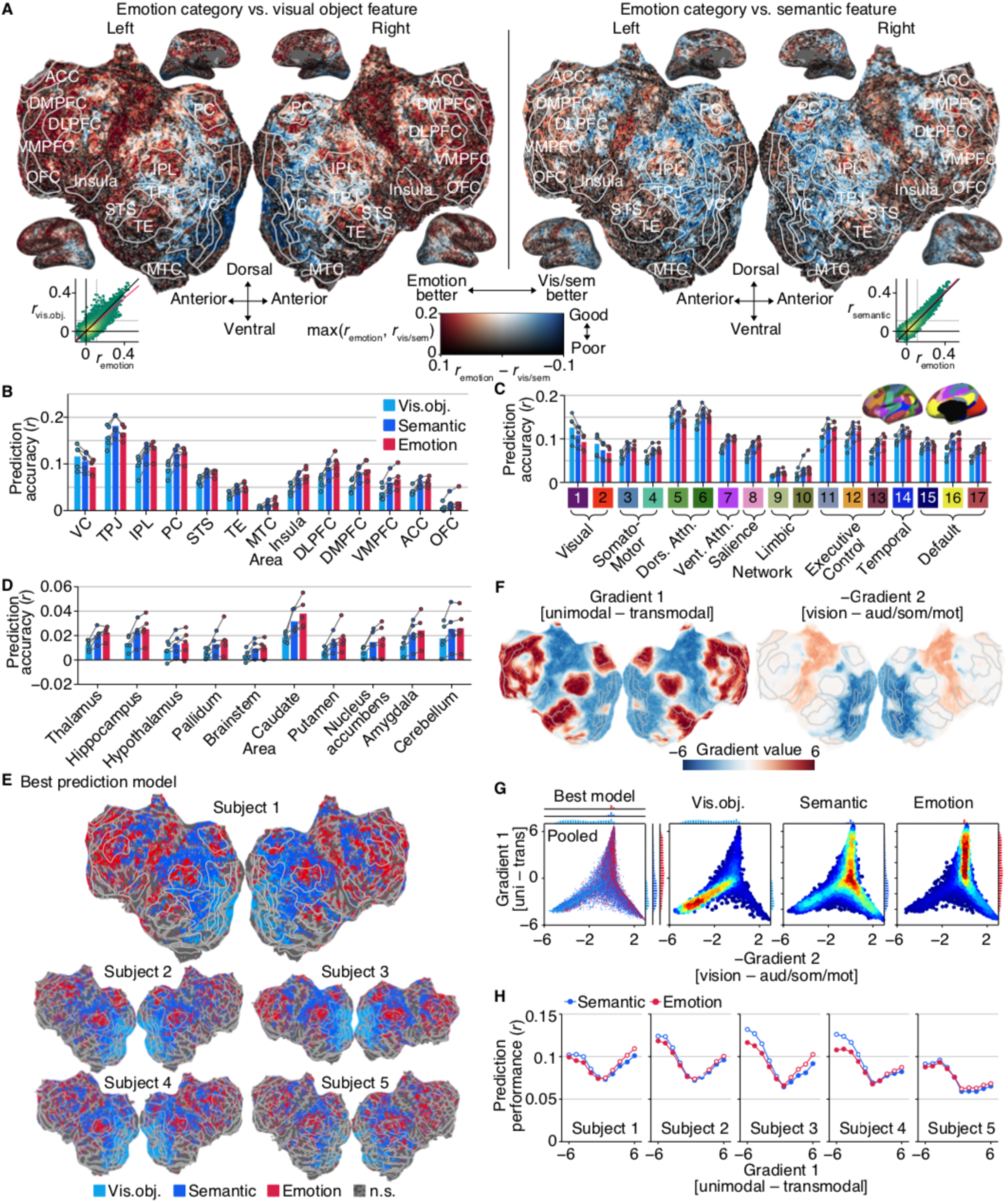
Disentangling emotional, visual object, and semantic feature encoding. (A) Differences in prediction accuracies of emotion, visual object, and semantic models (Subject 1, see Figure S4A for the other subjects). Inserted scatter plots follow the same conventions as Figure 4B. (B) Mean prediction accuracies in individual brain regions. Dots indicate accuracies of individual subjects. (C) Mean prediction accuracies in individual global networks. Inset offers the definitions of individual networks (see Figure S4C for a larger view). Conventions are the same as (B). (D) Mean prediction accuracies in individual subcortical regions. Conventions are the same as (B). (E) Best models among emotion (category), visual object, and semantic models (see Figure S4D for results with the dimension model). (F) Gradient values of the first and second PG axes. The sign of second PG axis is negative, following the convention of Margulies et al. (2016). (G) Joint and marginal distributions of the best models in PG space (five subjects pooled, see Figure S4E for individual subjects). Voxels with significant accuracy from either the visual object, semantic, or emotion models are shown (permutation test, *p* < 0.01, Bonferroni correction by the number of voxels). (H) Mean prediction accuracies along the first PG axis. Voxels were assigned to ten bins, or levels of the first axis, such that each bin consists of a roughly equal number of voxels. White circles indicate significant performance over the opponent model (two-tailed paired t-test, *p* < 0.01, Bonferroni correction by the numbers of PG levels and subjects).

### Visual object, semantic, and emotion encoding models along the cortical hierarchy

To better understand which brain regions preferentially represent emotion, we drew on the concept of the “principal gradient (PG)”, which is estimated from functional connectivity patterns (Margulies et al., 2016) and describes global hierarchical gradients in cortical organization (Huntenburg et al., 2019). Margulies et al. (2016) used a dimensionality reduction method to explore principal components of variances in cortical connectivity patterns, and found that the first component differentiates between unimodal and transmodal brain regions (Figure 5F left), and the second component differentiates between brain regions that represent different modalities (from visual to auditory and somatomotor cortices, Figure 5F right). The meta-analysis by Margulies et al. (2016) and a review study by Huntenburg et al. (2019) suggest that different regions along the PGs are responsible for broadly different brain functions.

Here, we tested empirically whether the PGs could account for the spatial arrangement of voxels preferentially representing visual object, semantic, and emotion information using encoding performances of models constructed from those different features. We plotted voxels predicted with significant accuracy by the visual object, semantic, or emotion models within the two-dimensional space defined by their values along the first and second PGs (Figure 5G, five subjects pooled, see Figure S4E for individual subjects). Voxels best predicted by each model occupied different locations in the PG space. Most importantly, along the first PG axis, voxels best predicted by the visual object, semantic, and emotion models were each densely distributed in regions with low, medium, and high values, respectively (ANOVA, interaction between frequency and level of the first PG axis, *p* < 0.01). In terms of raw prediction accuracy, voxel activity was better predicted by both the emotion and semantic models at low and high values of the first PG than at the midpoint. However, at low/high values of the first PG, the semantic/emotion model outperformed the emotion/semantic model, respectively (Figure 5H; ANOVA, interaction between accuracy and level of the first PG axis, *p* < 0.01 for all subjects). Because emotional responses are correlated with semantic features high in emotional affordance (e.g., injury, sexual activity), any brain regions predicted better by the emotion model than by the semantic model, even marginally, should be considered to encode emotional experience (or affordances for specific emotional experience). Hence, the present findings suggest that brain regions high along the first gradient axis — that is, transmodal regions — are central to the brain’s representation of emotion.

It is important to note that the results presented thus far were replicated using alternative analytic approaches, including the decoding (Figures S5A and B) and representational similarity analyses (Figures S5C and D; Kriegeskorte, et al., 2008). These analyses showed higher identification decoding accuracies (cf., Figure 3A) and representational similarities across the brain for the category than the dimension model (Figures S5A and C). In both cases, a highly similar set of brain regions was better fit by emotion models than by visual object model or semantic model (Figures S5B and D). These analyses demonstrate that our results establishing the primacy of emotion categories in neural representation are highly robust to alternative methodological choices.

Taken together, these results characterize how regions representing emotion are spatially arranged within the brain. While emotion-related representations are broadly distributed across the cortex, they are centered in regions that lie on the far end of a hierarchical gradient ranging from unimodal to transmodal regions. The representational shift along this gradient from visual to semantic to emotion-related representations may speak to the high levels of information integration and feature abstraction necessary for an emotional response.

### Distribution of brain activity patterns induced by emotional videos

While the above analyses addressed how particular dimensions, or kinds, of emotion are represented in the brain, the results thus far are not sufficient to reveal the distribution of neural responses to emotional stimuli along these dimensions, that is, how different kinds of emotion are distributed within a high dimensional space (Cowen and Keltner, 2017). To address the distribution of emotion-related brain activity, we performed unsupervised modeling using nonlinear dimensionality reduction and clustering analyses.

We first visually inspected the distribution of brain activity patterns induced by the 2181 emotional videos using the UMAP algorithm that we previously used to visualize the category emotion scores (cf., Figure 3D). In doing so, we selected voxels best predicted by either of the category or dimension encoding models with significant accuracy (cf., Figure S4D, including voxels in both cortical and subcortical regions), to focus on activity primarily representing emotion-related information. We then used activity patterns of those selected voxels as inputs to the UMAP algorithm, thus projecting a total of 2181 brain activity patterns into a two-dimensional space (see Transparent Methods: “Dimensionality reduction analysis”). The resulting maps colored based on ratings of the videos in terms of 27 emotion categories reveal a number of brain activity clusters that correspond to experiences of specific emotions (Figures 6A; five subjects averaged; e.g., sexual desire, disgust, and aesthetic appreciation), which were also observed even in maps of individual subjects. By contrast, maps colored by three representative affective dimensions (separately for positive and negative parts after 5 [neutral] was subtracted) do not highlight many distinct clusters (Figure 6B), indicating that the distribution of brain activity patterns was better captured by emotion categories.

**Figure 6.**
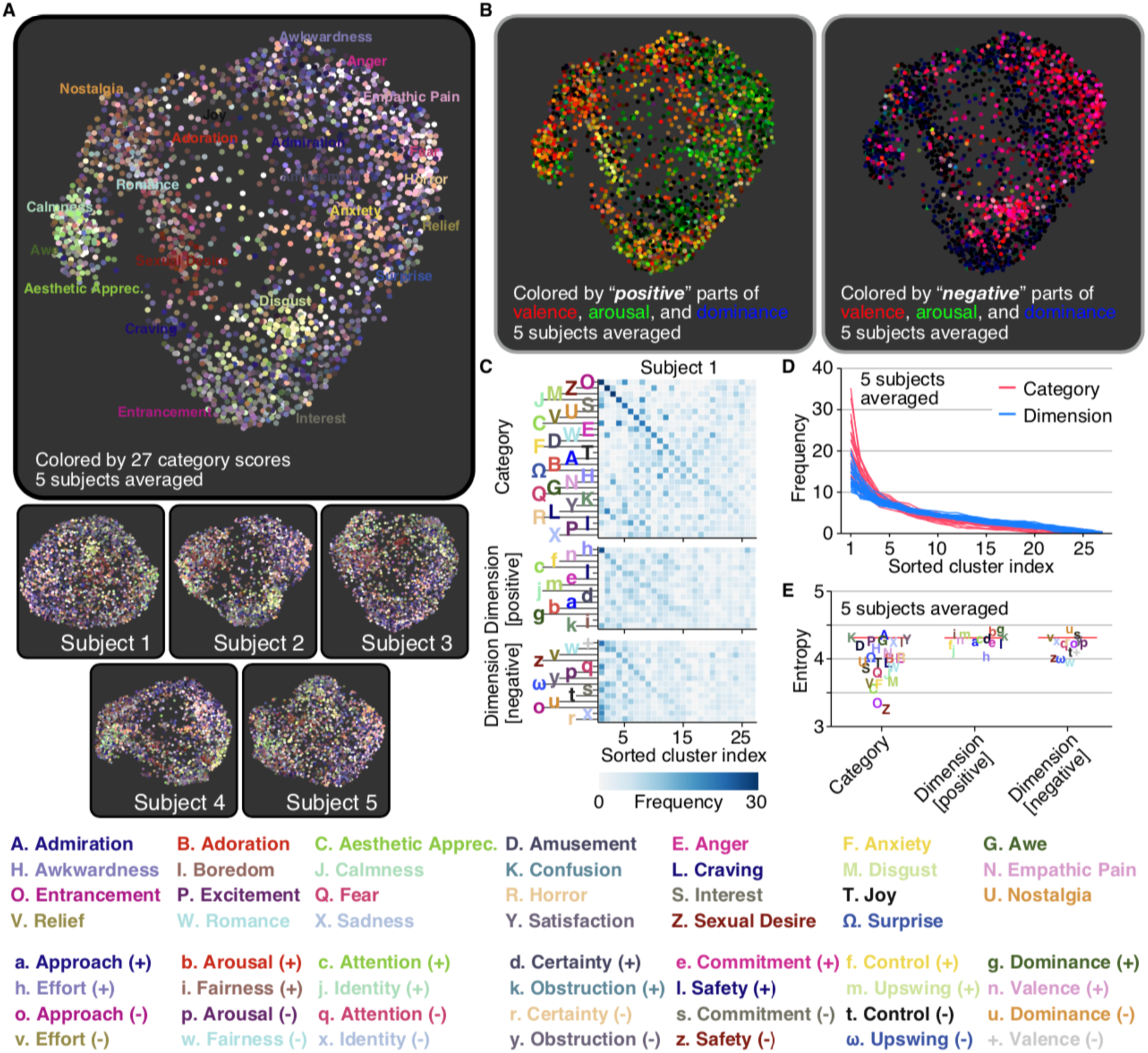
Distribution of brain activity patterns induced by emotional videos. (A) Maps of emotional experiences constructed from brain activity patterns. Each dot corresponds to each video. The same color schema as Figure 3D is used. (B) Maps of emotional experiences colored by positive or negative parts of valence, arousal, and dominance scores. (C) Distributions of top 5% high score samples of each emotion on 27 clusters derived from the brain (Subject 1, see Figure S6A for the other subjects). Clusters were separately sorted for the sets of the categories and positive/negative dimensions. (D) Sorted histograms for individual emotions. Lines indicate sorted histograms of 27 categories or 28 positive/negative dimensions (five subjects averaged, see Figures S6B and D for individual subjects and for results with different numbers of clusters). (E) Entropy of the top 5% high score sample distributions for each emotion (five subjects averaged; red lines, baseline, permutation test, *p* < 0.01, Bonferroni correction by the number of emotions; see Figures S6C and E for individual subjects and for results with different numbers of clusters).

We next quantified the degree to which brain activity patterns were distributed into cluster-like formations associated with specific emotions. For each subject, we applied k-means clustering to the 2181 brain activity patterns of the voxels selected as above, without any reference to the emotion scores of the eliciting videos. We set the number of clusters to 27, given that the previous study revealed 27 distinct varieties of emotional experiences elicited using the same videos (Cowen and Keltner, 2017), and we would expect at most one cluster per distinct emotion. Note, however, that similar results were obtained using different numbers of clusters (Figure S6). To characterize the emotional properties of each cluster, we examined how brain activity patterns corresponding to videos with top 5% high scores (109 videos) for each emotion are assigned to these clusters using scores of the previously reported 27 distinct categories and the positive and negative parts of affective dimensions (absolute values after 5 [neutral] was subtracted).

Histograms of the samples assigned to the 27 clusters for each emotion are shown in Figure 6C (see Figure S6A for the other subjects), revealing an important disparity between the categories and dimensions. Namely, top samples of several categories, including entrancement, sexual desire, and disgust, tended to belong to a small number of clusters, while those of affective dimensions tended to be broadly assigned to many clusters. This difference was quantified in two ways. First, the histograms of each emotion — independently sorted according to their frequencies within each cluster — were steeper for the categories than dimensions (Figure 6D; five subjects averaged; ANOVA, interaction between frequency and sorted clusters, *p* < 0.01; see Figure S6B for individual subjects). Second, entropy calculated for each histogram of individual emotions tended to be low for many categories (e.g., O: entrancement, Z: sexual desire, and C: aesthetic appreciation) compared to most of the dimensions (Figure 6E, five subjects averaged, see Figure S6C for individual subjects).

Together, these results indicate that brain activity patterns induced by emotional videos have cluster-like distributions that align to a greater degree with categories than with dimensions. Meanwhile, close inspection of the map (Figure 6A) also reveals that the patterns of brain activity corresponding to certain emotions (e.g., fear and horror) seem to be not entirely discrete but overlapping in their distribution. Indeed, top samples of several categories (e.g., W: romance and Z: sexual desire; C: aesthetic appreciation and J: calmness) tended to fall into overlapping clusters (Figure 6C and Figure S6A). This finding may shed light on the biological basis of gradients between distinct emotional experiences, originally revealed in self-report (Cowen and Keltner, 2017), by suggesting that they correspond to gradients in evoked patterns of brain activity.

## Discussion

We have sought to characterize the neural substrates of emotional experience by analyzing activity induced by the richest set of emotional visual stimuli investigated to date in neuroscience — 2181 emotionally evocative videos annotated with 34 emotion categories and 14 affective dimensions — using neural decoding, voxel-wise encoding modeling, and unsupervised modeling methods. We found that ratings of individual emotions could accurately be predicted from activity patterns in many brain regions, revealing that distributed brain networks contributed in distinct ways to the representation of individual emotions in a highly consistent manner across subjects. Using voxel-wise encoding models constructed from the category and dimension ratings, we found that the brain representation of emotion categories explained greater variability in brain activity than that of affective dimensions in almost all brain regions. By comparing encoding performances between visual object, semantic, and emotion models, we confirmed that responses in a variety of brain regions were explained by features related to emotion rather than visual and semantic covariates. Those emotion-related representations were broadly distributed, but commonly situated at the far end of a cortical gradient that runs from unimodal to transmodal regions. Analyses using unsupervised modeling revealed that the distribution of brain activity patterns evoked by emotional videos was structured with clusters associated with particular emotion categories bridged by gradients between related emotions. Taken together, our results demonstrate that neural representations of diverse emotional experiences during video viewing are high-dimensional, categorical, and distributed across transmodal brain regions.

Our decoding analysis revealed that brain regions that contributed to the representation of individual emotions were distinct for different emotions and similar across individuals (Figure 2). Emotions were represented not by simple one-to-one mappings between particular emotions and brain regions (e.g., fear and the amygdala) but by more complex configurations across multiple networks, consistent with notions of distributed, network-based representations of specific emotions (Hamaan, 2012; Lindquist and Barrett, 2012; Saarimäki et al., 2016). In previous studies, emotion representations were found largely in transmodal brain regions, which are activated by tasks that rely upon social cognition, autobiographical memory, reward-based decision making, and other higher cognitive functions (Margulies et al., 2016). One exception from our analysis is that somatosensory regions were also found to encode empathic pain (Figure 2), consistent with notions that our empathy for others’ injuries arises in part via mental simulation (Bufalari et al., 2007). Overall, these findings may reflect how experiences of individual emotions involve multiple interacting systems that automatically enter distinct states in response to significant threats and opportunities in the environment (Cowen, 2019; Seo et al., 2019). Given that these systems encode emotional experience (or affordances) even during passive viewing of videos, these findings support the view that emotions rely on system-wide adaptations that proactively recruit psychological faculties likely to support adaptive behavioral responses even before a decision is made to respond.

Our findings reconcile a history of observations that localized brain lesions can have highly specific effects on emotional behavior (e.g., Buchanan et al., 2004; Calder et al., 2000; Ciaramelli et al., 2013; Kim et al., 1993; Scott et al., 1997) with findings that these behaviors lack simple one-to-one mappings onto activity in localized brain regions (Lindquist and Barrett, 2012). If a given emotion is represented in a specific mode of activity within a network of interacting brain regions, damage to various hubs within this network would have the potential to disrupt specific behaviors associated with that emotion in differential ways (a premise further supported by findings from systems neuroscience in animal models, e.g., Kim et al., 1993; Seo et al., 2019). Our findings reveal how a set of transmodal brain regions centered in the default mode network undergo different modes of activity for different emotions. Intriguingly, the default mode network can in this way be considered to have not just one mode, but many modes of activity corresponding to different emotions, with varying subregions of the network being differentially involved in each emotion (Figure 2).

Whether the neural encoding of specific emotion categories (e.g., “anger”) can be reduced to the encoding of few broad affective dimensions such as valence, arousal, and certainty is an active area of debate that informs theories of emotion (Lindquist and Barrett, 2012; Kragel et al., 2019; Saarimäki et al., 2018). Our encoding and decoding analyses demonstrate that emotion categories reliably capture neural representations that are more specific than those captured by a wide range of affective dimensions throughout the brain (Figure 4). These results converge with recent behavioral studies revealing how emotion categories organize reported emotional experiences and emotion recognition from nonverbal signaling behavior (Cowen et al., 2019a; Cowen and Keltner, 2017). Previous studies have pitted categorical and dimensional emotion models against one another (Hamaan, 2012), but done so with only a few emotions (e.g., happiness, anger, fear, sadness, disgust, and surprise) or two dimensions (valence, arousal), conflating distinct emotional experiences within and across studies. In the current study, we drew upon recent advances in the range of emotions considered to be distinct and fitted models accounting for a fuller array of appraisal dimensions of theoretical relevance to enable a more robust and thorough comparison of categorical and dimensional models of emotion representation. Our results provide strong support for a categorical approach to emotional experience, or theories that propose neural adaptations for a large set of appraisal dimensions specific enough to account for dozens of distinct emotion categories (e.g., Skerry and Saxe, 2015), models of which would consequently bear the same information as our category model (although further work would be needed to establish such a comprehensive set of appraisals; see Cowen et al., 2019b for discussion). Additionally, using unsupervised modeling methods, we found that the distribution of emotion-related brain activity patterns in regions encoding emotion had cluster-like formations corresponding to particular emotion categories but not broad appraisals (Figure 6A), further supporting the conceptualization of the emotional response in terms of self-reported experiences such as “anxiety”, “disgust”, and “entrancement”. Taken together, these results point to a high-dimensional, categorical space in the neural representation of emotion.

As done in previous studies using voxel-wise modeling in specific sensory regions (de Heer et al., 2017; Güçlü and van Gerven, 2015; Kell et al., 2018; Lescroart and Gallant, 2019), we compared performances of encoding models constructed from different features (emotion scores, visual object features, and semantic features), and uncovered a distributed set of brain regions that encode emotional experience (Figure 5). The spatial arrangement of these emotion-related brain regions overlapped with that of the default mode network (Figure 5C) and corresponded to the far end of a gradient from unimodal to transmodal regions (Figures 5G and H) that are located at geodesically distant points from primary sensory cortices (Margulies et al., 2016). Our analysis in a single study revealed a gradual representational shift along the unimodal-transmodal gradient from visual to semantic to emotion information, perhaps reflecting a global hierarchy of integration and feature abstraction from sensory inputs (Huntenburg et al., 2017).

One issue worth noting is that, because each stimulus was presented only once, we were unable to estimate the proportion of systematic variance in voxel responses that was left unexplained by our models. Many voxel-wise encoding modeling studies have relied on repeated stimulus presentations to determine the upper bound of model prediction accuracies (Huth et al., 2016; Kell et al., 2018; Lescroart and Gallant, 2019). However, responses to emotional stimuli are likely to change with repeated exposure to the stimuli, given that many emotional responses (e.g., surprise) can be affected by past exposure and expectations. For this reason, we performed our analyses on data from single trials. Nevertheless, the model prediction accuracies we observed were moderately high by fMRI standards. This might be attributable in part to the potency of naturalistic stimuli in evoking widespread brain responses (Hamilton and Huth, 2018), and, more specifically, to the emotional intensity of many of the video stimuli we used (Cowen and Keltner, 2017).

On this point, it is worth considering how investigations of brain representations of emotion may depend on the richness of the experimental stimuli used. The 2181 videos studied here captured a wide spectrum of psychologically significant contexts. By contrast, most previous work has investigated neural underpinnings of emotion using a far narrower range of eliciting stimuli and a small set of emotions (Barrett, 2017; Celeghin et al., 2017; Giordano et al., 2018; Hamaan, 2012; Kragel and LaBar, 2015; Lindquist and Barrett, 2012; Saarimäki et al., 2016; Satpute and Lindquist, 2019; Wager et al., 2015). Only recently have several studies investigated neural representations of emotion with richer models of emotion, but they have still relied on a relatively narrow range of stimuli (Koide-Majima, et al., 2018; Kragel et al., 2019; Saarimäki et al., 2018; Skerry and Saxe, 2015). In assembling the present stimulus set, Cowen and Keltner (2017) drew on the recent availability of highly evocative stimuli on the internet to capture more extensive, diverse, and poignant stimuli, although it is worth noting that these stimuli are still only visual in nature. Future studies could build on this work by including auditory content, content with personal significance to the subject (e.g., imagery of loved ones), or tasks that include behavioral involvement of the subject (e.g., games, social interaction). As the field of affective science moves forward, the present study reveals how a more extensive analysis of whole-brain responses to a much richer and more extensive set of experimental conditions, open to diverse analytical approaches, can yield more definitive results.

This approach allowed rich and nuanced emotion categories to be accurately predicted from brain activity patterns (Figures 2A and B) and even made it possible to accurately identify eliciting stimuli based on patterns of decoded emotion scores (Figure 3). Previous decoding studies of emotion have relied on discrete classification into a small set of emotion categories (Kragel et al., 2016; Kragel and LaBar, 2015; Saarimäki et al., 2016; Saarimäki et al., 2018; Sitaram et al., 2011), which may conflate representations of diverse emotional states grouped into each category (Clark-Polner et al., 2016). By predicting continuous intensities of many distinct emotions simultaneously, a more comprehensive picture of emotional states can be characterized. We visualized these nuanced emotional states by combining decoded scores of richly varying emotion categories with a UMAP algorithm (Figure 3D). This approach can provide a tool for various applications, such as product evaluation in neuro-marketing (Nishida and Nishimoto, 2018). Furthermore, in light of recent findings of a broad overlap between neural representations of perceived and internally-generated experiences (Horikawa et al., 2013; Kragel et al., 2016; Tusche et al., 2014), decoders trained for perceived emotions may generalize to emotional states that occur spontaneously during dreams and mind wandering, with potential clinical applications (Kragel et al., 2016; Sitaram et al., 2011).

A critical question in neuroscience is where emotions evoked by different modalities of sensory input are encoded in the brain (Chikazoe et al., 2014; Kragel et al., 2016; Skerry and Saxe, 2014). Although here we used only visual stimuli (with no sound), we found that emotion-related representations were not primarily found in visual cortex (Kragel et al., 2019) but in many transmodal and frontal regions that have been implicated in representing other modalities of emotionally evocative stimuli and tasks, including verbal stimuli (Huth et al., 2016; Skerry and Saxe, 2015), visual and gustatory stimuli (Chikazoe et al., 2014), and volitional and spontaneous imagery (Tusche et al., 2014). A few studies have investigated commonalities in the neural representation of emotion across multiple stimulus modalities, but have focused on narrower ranges of stimuli and emotions (e.g., basic emotions: Peelen et al., 2010; valence: Chikazoe et al., 2014; Skerry and Saxe, 2015). As behavioral studies generate richer and diverse experimental resources for investigating multiple modalities of emotions (e.g., resources for emotional responses to facial expression, speech prosody, nonverbal vocalization, music, games, and social interaction; Cowen and Keltner, 2019; Cowen et al., 2018, 2019a, 2020), cross-modal neural representations of emotions should be investigated more deeply with richer models of emotion.

## Limitations of the Study

One limitation of the present study was that the emotion ratings we used were averages from multiple raters and not from the subjects of our fMRI experiment. While our decoding and encoding models were able to establish systematic statistical relationships between those ratings and the brain activity patterns of our subjects, the reported feelings of those raters may not completely match those of our subjects. Hence, to the extent that neural representations of emotion are subject to individual differences in culture, demographic variables (gender, age), and personality, future studies might uncover more robust neural representations by incorporating these variables.

Another potential concern regarding the present investigation is the confounding of first-person and third-person emotions. The brain regions in which the category model outperformed the visual object and semantic models (e.g., IPL, VMPFC) overlapped with brain regions that have been implicated in a “theory of mind network” (Skerry and Saxe, 2015). It is possible that the neural representations uncovered here reflect information not only regarding first-person emotional experiences but also regarding predictions of how others might feel. While the observed overlap may also indicate common neural substrates for first- and third-person emotions, future experiments are needed to dissociate these phenomena using more carefully designed experiments, perhaps including psychophysiological measures indicative of first-person emotional experience.

Finally, as we allowed the subjects to freely view presented video stimuli, differential gaze patterns for each emotional experience might have some influence on our brain data. While the neural representation of gaze is known to be highly localized in specific brain regions (Connolly et al., 2005; Sestieri et al., 2008; Culham et al., 1998), modeling effects of emotion on gaze patterns would help to more precisely characterize neural representations associated with subjective feelings of emotions.

## Acknowledgments

The authors thank Mitsuaki Tsukamoto, Hiroaki Yamane, and Shuntaro C. Aoki for help with data collection and software setups. This study was conducted using the MRI scanner and related facilities of Kokoro Research Center, Kyoto University. This research was supported by grants from the New Energy and Industrial Technology Development Organization (NEDO), JSPS KAKENHI Grant number JP15H05710, JP15H05920, and JP17K12771, JST CREST Grant Number JPMJCR18A5, and JST PRESTO Grant Number JPMJPR185B Japan.

## Author Contributions

Conceptualization, T.H. and Y.K.; Methodology, T.H.; Validation, T.H.; Formal Analysis, T.H.; Investigation, T.H.; Resources, Y.K.; Writing – Original Draft, T.H.; Writing – Review & Editing, T.H., A.S.C., D.K., and Y.K.; Visualization, T.H.; Supervision, Y.K.; Funding Acquisition, T.H. and Y.K.

