## Supplemental Information for "The neural representation of visually evoked emotion is high-dimensional, categorical, and distributed across transmodal brain regions"

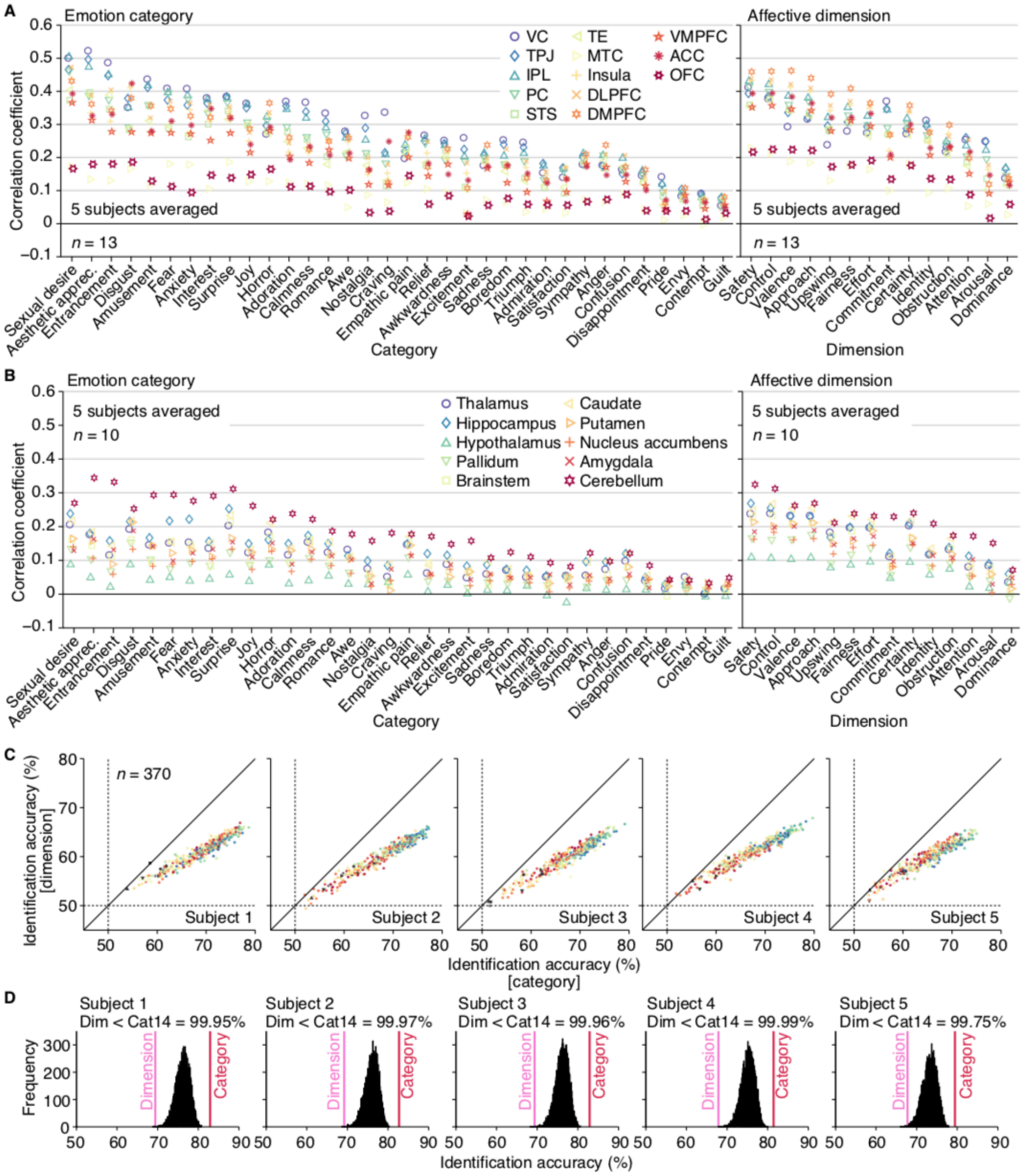


**Figure S1. Performances of decoding analysis for emotion scores. (Related to Figures 2 and 3)**

(A) Decoding accuracy for individual emotions predicted from activities in representative cortical regions. The decoding analysis of individual emotion scores (cf., Figures 2A and B) was performed from brain activity patterns in several cortical regions (see Transparent Methods: “Regions of interest (ROI)” for definitions of individual cortical regions). Dots indicate accuracies obtained from individual cortical regions (five subjects averaged).

(B) Decoding accuracy for individual emotions predicted from activities in subcortical regions. Conventions are the same as (A).

(C) Mean video identification accuracies from region-wise decoders of individual subjects. Conventions are the same as Figure 3A.

(C) Distributions of video identification accuracies obtained from randomly selected 14 emotion category scores. The video identification analysis by ensemble decoders (cf., Figure 3B) was performed for 10,000 times while randomly selecting different combinations of 14 emotion category scores from the original 34 emotion category scores. The identification accuracies obtained with this procedure were compared with the accuracy from the 14 affective dimensions. The estimated accuracies from more than 99% of 14 randomly selected emotion categories outperformed the accuracy from the 14 affective dimensions. The results suggest that the superiority of the 34 emotion categories over the 14 affective dimensions (Figure 3B) were not solely due to the differences of the number of emotions used for identification.

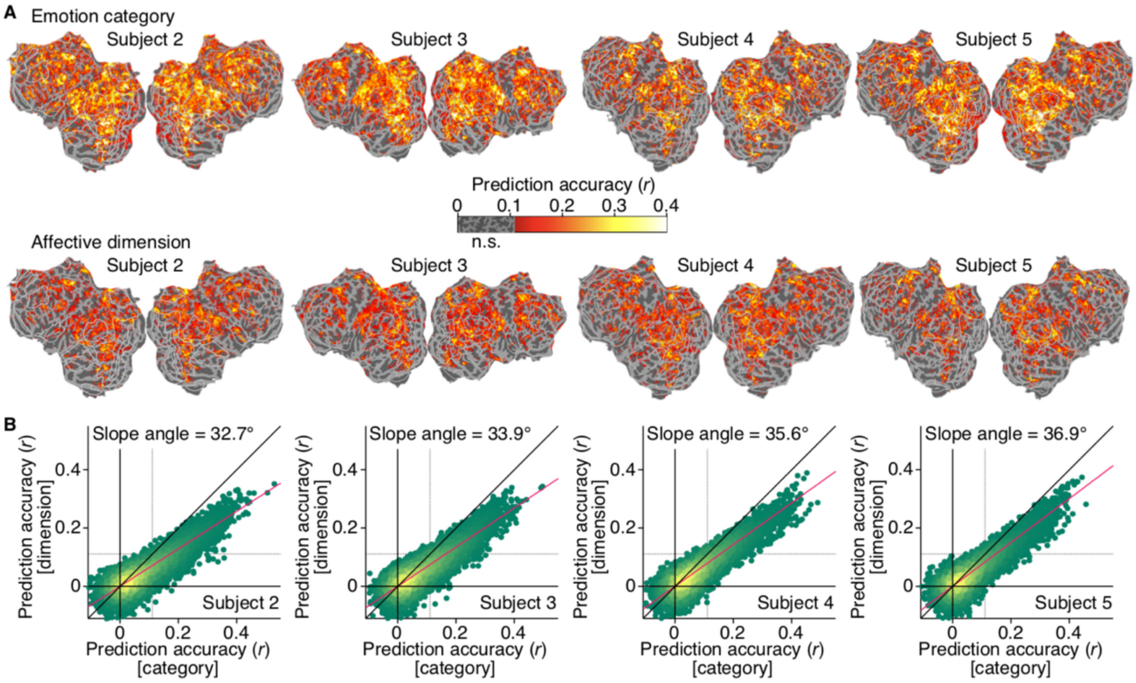


**Figure S2. Performance of encoding models constructed from emotional scores for individual subjects. (Related to Figure 4)**

(A) Prediction accuracies of emotion encoding models. Conventions are the same with Figure 4A.

(B) Prediction accuracies o individual voxels. Conventions are the same with Figure 4B.

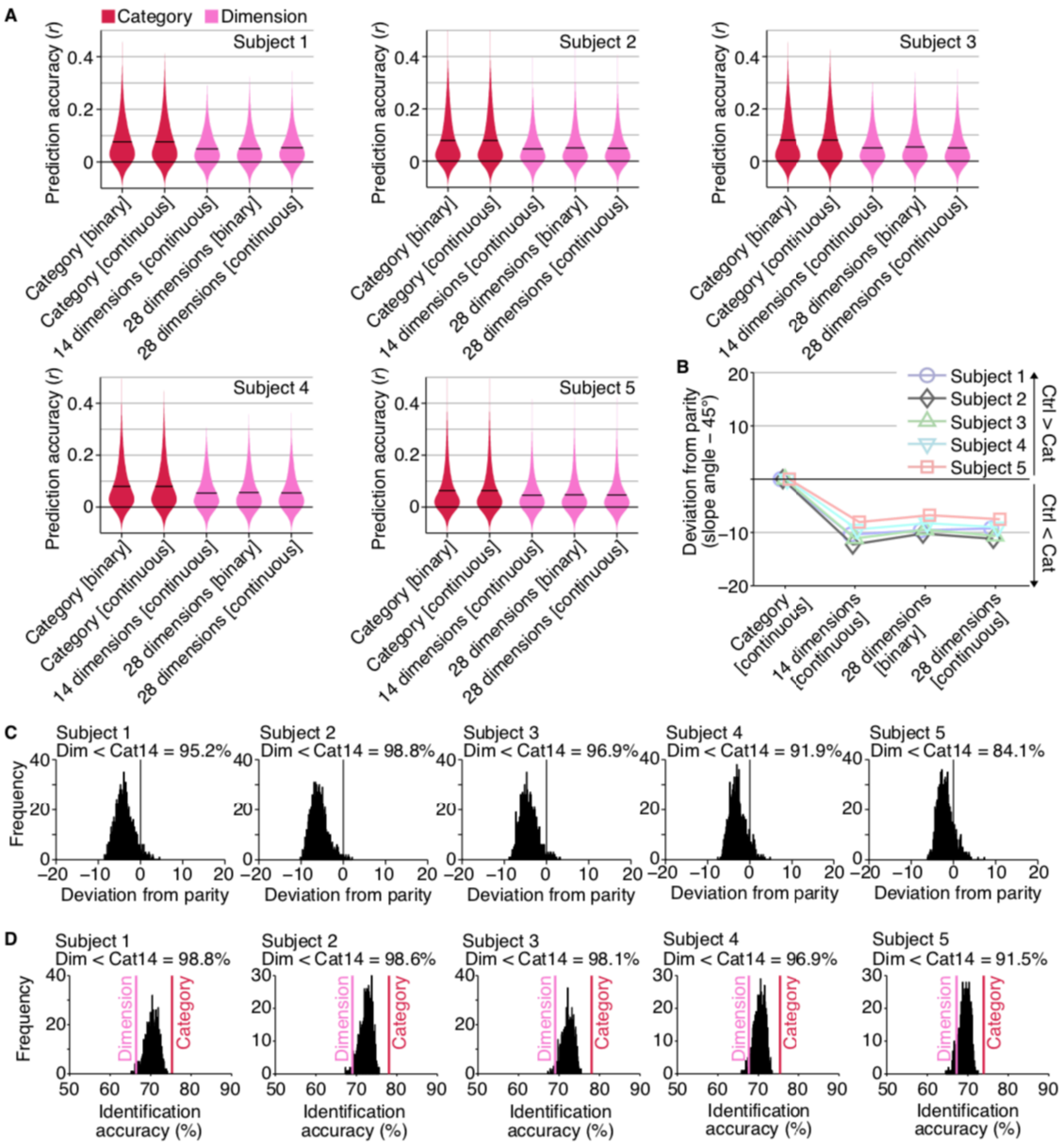


**Figure S3. Control analyses for the performance comparisons between category and dimension encoding models. (Related to Figure 4)**

(A) Distributions of prediction accuracies of all voxels from multiple variants of category models and dimension models. Because the methods for collecting scores of emotion categories and affective dimensions were different (see Transparent Methods: “Video stimulus labeling”), differences of encoding performances might be attributable to such differences. To compensate for the differences, we constructed encoding models from different versions of emotion category and affective dimension scores used in the main analyses, and compared encoding model performances from those multiple variants. For the emotion category scores, we have used binarized scores reported by individual human raters (originally ranged between 0 to 100) in the

main analysis (“category [binary]”; cf., Figure 4). We here also tested another type of emotion category scores without binarization (mean of reported scores [ranged between 0 to 100], “category [continuous]”). In either case, high and low values of the emotion category scores can be interpreted whether that emotions exist or not. On the other hand, because scores of the affective dimensions were collected with 9-scale Likert scale (ranged between 1 to 9, “14 dimensions [continuous]”), the high and low values should be interpreted differently (e.g., positive and negative), and low values should be interpreted as strong negative emotions rather than no emotion. To compensate for the difference from the category scores, we tested other types of affective dimension scores by first subtracting 5 [neutral] from original values (-4 to 4), taking both positive and negative values separately, and concatenating the positive (0 to 4) and negative (-4 to 0) parts to yield a total of 28 unipolar affective dimension scores (“28 dimensions [continuous]”). This score conversion was done for scores of individual raters and converted scores were averaged across multiple raters. Furthermore, similar to the emotion category scores, we also tested the binarized version of affective dimension scores by binarizing zero or non-zero values of the continuous 28-dimensional scores of individual raters to zero/one values and averaging them across multiple raters (“28 dimensional [binary]”). These score conversions had little effect on encoding accuracies, showing higher mean encoding accuracies for the emotion category models than the affective dimension models.
(B) Comparisons of prediction accuracies between the emotion category models and the other control models. The encoding performance obtained from the original emotion category model was directly compared with the performances from models constructed with the other variants of emotion scores (cf., Figure S3A) based on slope angles of best linear fit estimated from encoding accuracies of all voxels across the whole brain. The results showed equivalent performances between the two category models, whereas the original category model outperformed all affective dimension models, suggesting that the differences of the data collection methods were not main factor of the superiority of the emotion category model.
(C) Distributions of deviations of slope angles from the parity compared between the dimension models and category models constructed with randomly selected 14 emotion category scores. Encoding models were constructed from 14 randomly selected emotion categories for 1,000 times, and obtained encoding accuracies were each compared with the accuracy from the 14 affective dimensions based on slope angles of best linear fit estimated from encoding accuracies of all voxels across the whole brain. The results showed that on average more than 93.4% models constructed from 14 randomly selected emotion categories outperformed the accuracy from the 14 affective dimensions (five subjects averaged).
(D) Distributions of video identification accuracies obtained from encoding models constructed from randomly selected 14 emotion categories. The video identification analysis (cf., Figure 4F) was performed with models constructed with 14 randomly selected emotion categories for 1,000 times (cf., Figure S3C). The identification accuracies obtained with this procedure were

compared with the accuracy from the 14 affective dimension model (Figure 4F). The results showed that on average more than 96.8% models obtained from 14 randomly selected emotion categories outperformed the accuracy from the 14 affective dimensions (five subjects averaged). Taken together with (C), the results suggest that the superiority of the 34 emotion categories over the 14 affective dimensions were not solely due to the differences of the number of emotions used for the encoding analysis.

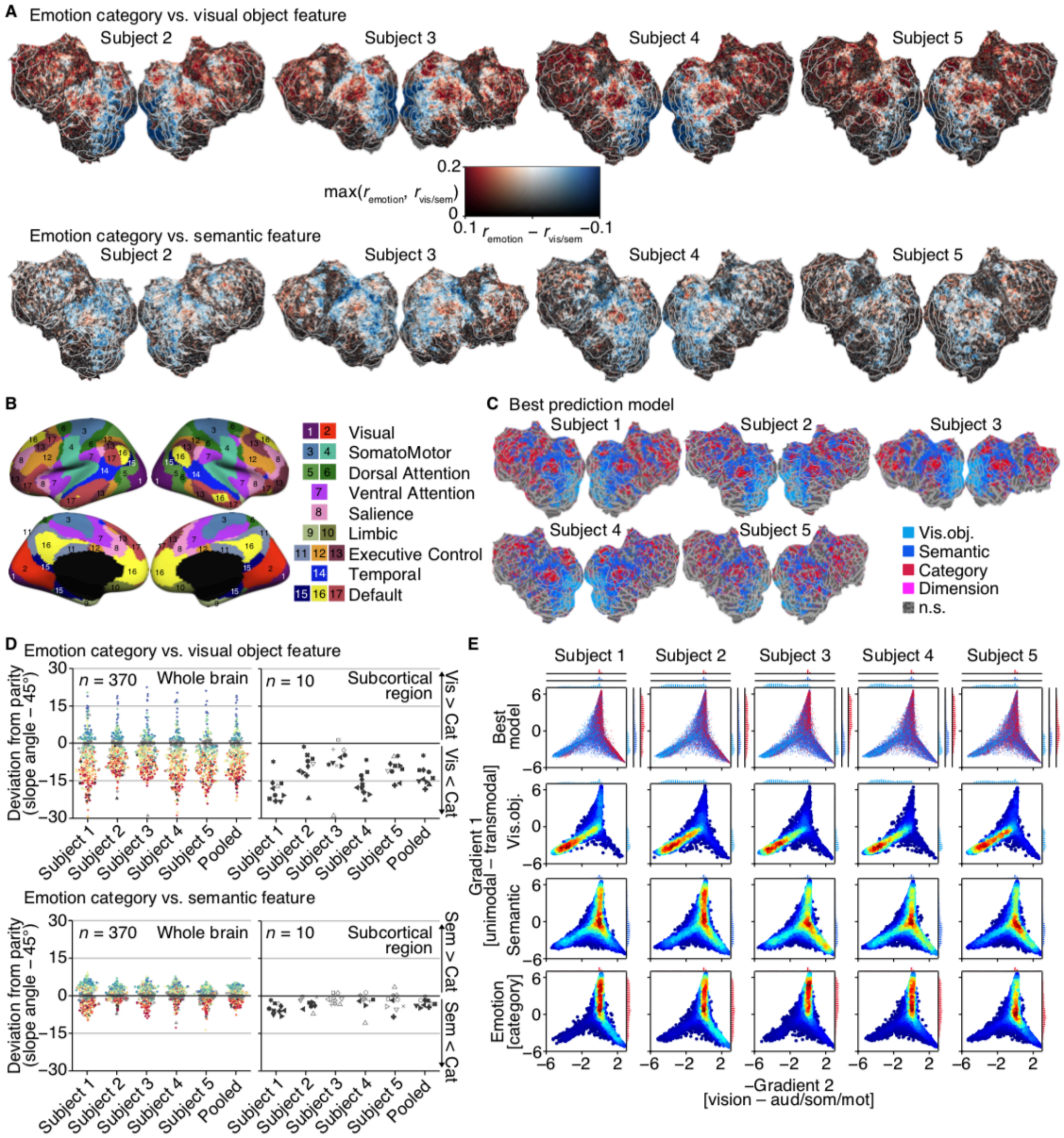


**Figure S4. Comparisons of encoding accuracy based on emotion, visual object, and semantic models for individual subjects. (Related to Figure 5)**

(A) Differences in prediction accuracies of emotion, visual object, and semantic models.

(B) Definition of global networks (see Yeo et al., 2011 for details).

(C) Best models among visual object, semantic, category, and dimension models.

(D) Comparisons of prediction accuracies for individual brain regions. Conventions are the same as Figure 4D. To examine the similarity of the distributions of emotion-related regions across subjects, the Pearson correlation coefficients were calculated between the estimated slope angles of all brain regions ($n$ = 370) from each pair of subjects ($n$ = 10). The analysis showed highly positive correlations for both comparisons between the emotion and visual object models ($r$ = 0.843, averaged across pairs) and between the emotion and semantic models ($r$ =

0.624, averaged across pairs), suggesting that the distributions of emotion-related brain regions were similar across subjects.

(E) Joint and marginal distributions of the best models in principal gradient space for individual subjects. Conventions are the same as Figure 5G.

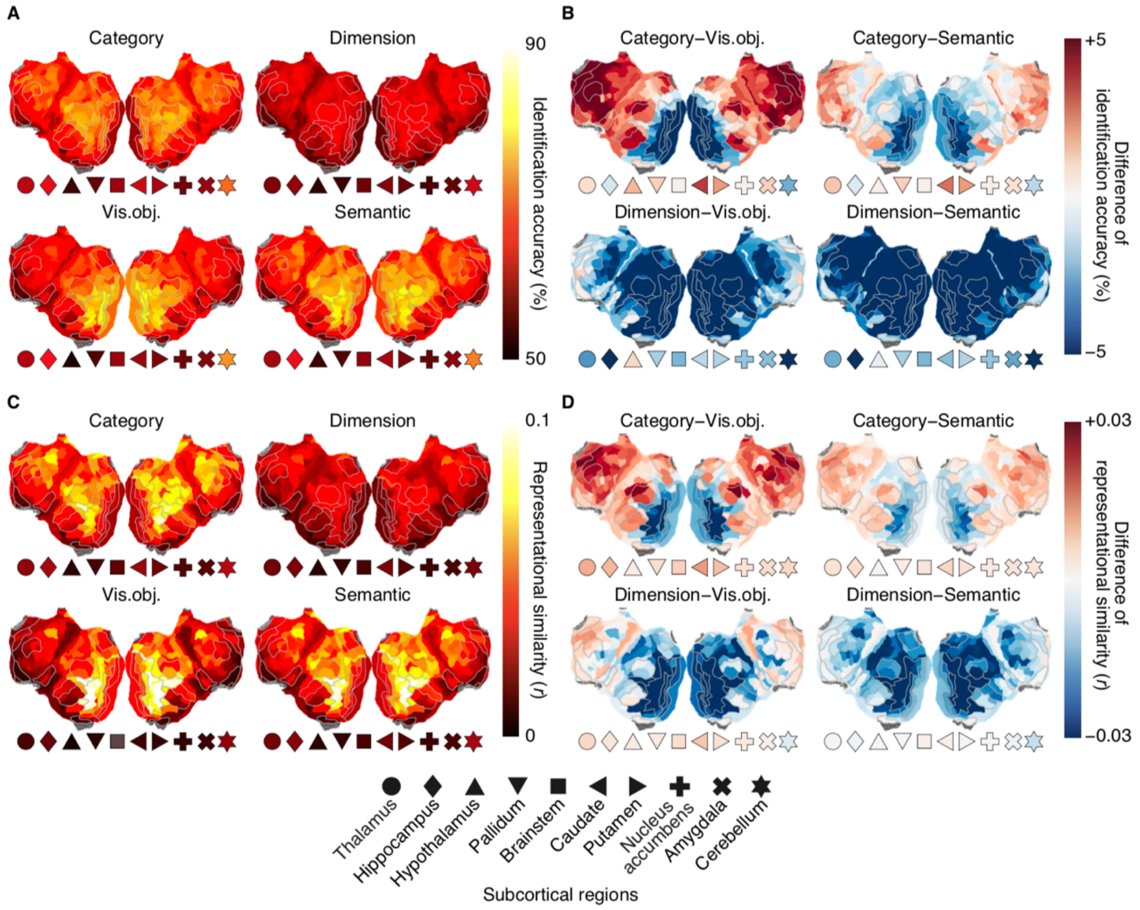


**Figure S5. Performances of decoding analysis and representational similarity analysis across the whole cortex. (Related to Figure 5)**

(A) Video identification accuracies via decoding analysis of individual features (five subjects averaged). The video identification analysis performed using decoded emotional scores (cf., Figures 3A and S1C) was also performed with visual object features and semantic features via decoding analysis of individual features. Mean accuracies of the identification analysis via decoded scores/features from five subjects (cf., Figure S1C) were averaged for each brain region, and were projected onto the cortical surface of Subject 1.

(B) Differences in video identification accuracies via decoding analysis between two different features. Conventions are the same as (A).

(C) Representational similarities for individual feature set. The representational similarity analysis (Kriegeskorte et al., 2008) was performed to evaluate the similarity of representational similarity matrices (RSM) constructed from patterns of brain activities and scores/features (see Transparent Methods: "Representational similarity analysis" for details). For each set of features (e.g., 34 emotion category scores), correlation coefficients between two RSMs (one from the brain, and the other from scores/features) were calculated for individual brain regions ($n$ = 370). The estimated representational similarities for individual brain regions were averaged across five subjects, and were projected onto the cortical surface of Subject 1.

(D) Differences in representational similarities between two different feature sets. Conventions are the same as (C).

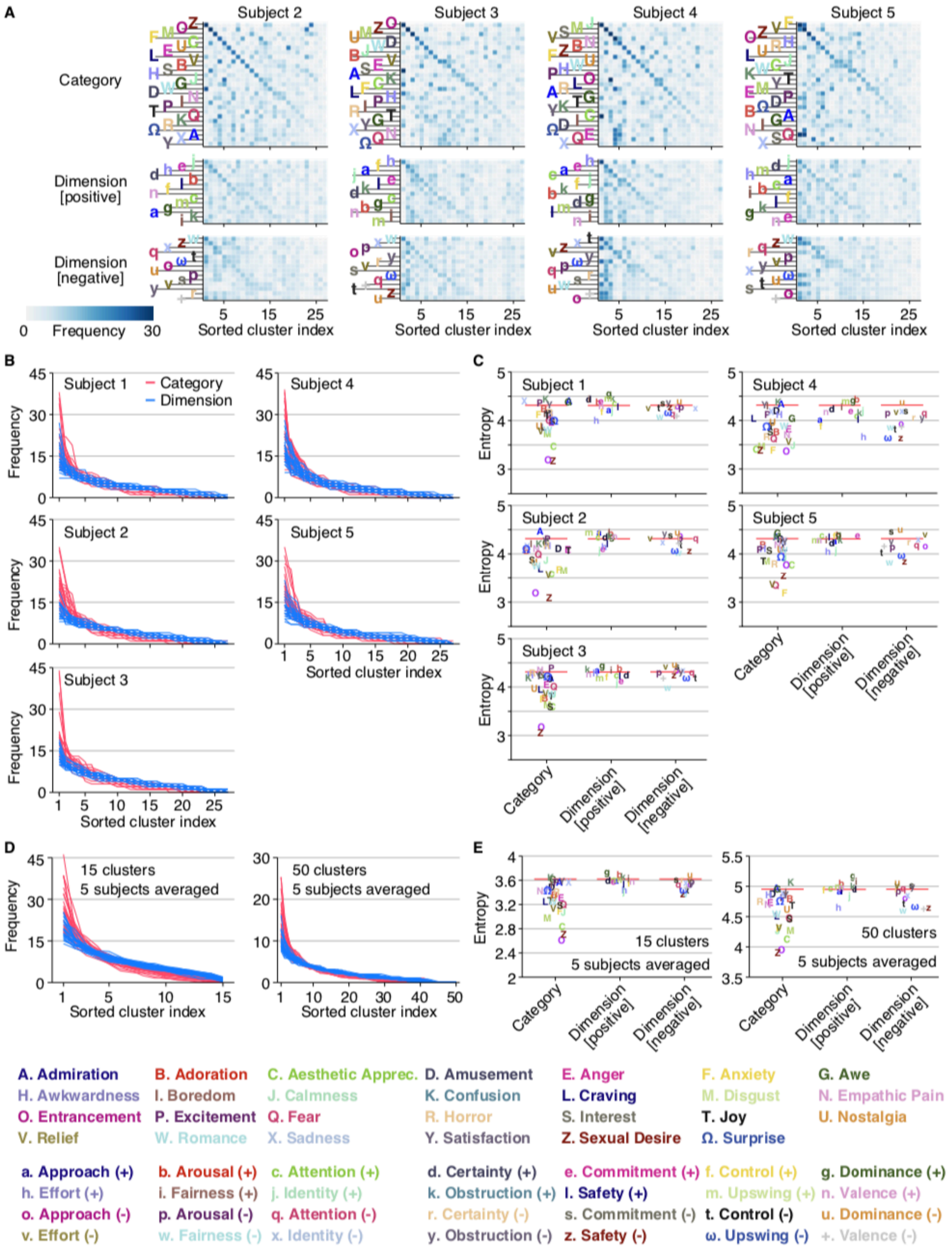


**Figure S6. Clustering analysis on brain activity patterns induced by emotion evocative short videos for individual subjects. (Related to Figure 6)**

(A) Distributions of top 5% high score samples of individual emotions on 27 clusters derived from brain activity patterns for individual subjects. Conventions are the same with Figure 6C.

(B) Sorted histograms of individual emotions for individual subjects. Conventions are the same

with Figure 6D.
(C) Entropy of the top 5% high score sample distributions of each emotion for individual subjects. Conventions are the same with Figure 6E.
(D) Sorted histograms of individual emotions obtained with 15 and 50 clusters. Conventions are the same with Figures 6D.
(E) Entropy of the top 5% high score sample distributions of each emotion obtained with 15 and 50 clusters. Conventions are the same with Figures 6E.

**Transparent Methods**

### CONTACT FOR REAGENT AND RESOURCE SHARING

### EXPERIMENTAL MODEL AND SUBJECT DETAILS

#### Human subjects

Five healthy subjects with normal or corrected-to-normal vision participated in our experiments: Subject 1 (male, age 34), Subject 2 (male, age 23), Subject 3 (female, age 23), Subject 4 (male, age 22), and Subject 5 (male, age 27). This sample size was chosen on the basis of previous fMRI studies with similar experimental designs (Horikawa and Kamitani, 2017; Huth et al., 2016). All subjects provided written informed consent for participation in the experiments, and the study protocol was approved by the Ethics Committee of ATR.

### METHOD DETAILS

#### Emotional movie stimuli

The stimuli consisted of sequences of emotionally evocative short videos. The videos were originally collected by Cowen and Keltner (2017). The video dataset consisted of a total of 2196 videos (downloaded at 13 September, 2017; https://goo.gl/forms/XErJw9sBeyuOyp5Q2) whose durations ranged from ~0.15 s to ~90 s. Each video was resized so that the width and height of videos were both within 12 degree (the original aspect ratio was preserved) and was visually presented at the center of the screen on gray background (no sound was delivered in our experiment, while some videos originally contained sounds).

#### Experimental design

All video stimuli were rear-projected onto a screen in the fMRI scanner bore using a luminance-calibrated liquid crystal display projector. To minimize head movements during fMRI scanning, subjects were required to fix their heads using a custom-molded bite-bar individually made for each subject except for the case where subjects were reluctant to use the bite-bar (a subset of sessions with Subject 5). Data from each subject were collected over multiple scanning sessions spanning approximately 2 months. On each experimental day, one consecutive session was conducted for a maximum of 2 hours. Subjects were given adequate time for rest between runs (every 7–10 min) and were allowed to take a break or stop the experiment at any time.

The video presentation experiment consisted of a total of 61 separate runs. Each run comprised 36 stimulus blocks whose durations differed across blocks depending on the durations of videos presented in each stimulus block. For stimulus blocks with videos shorter than 8 s, the same video stimulus was repeatedly presented until the total presentation duration went beyond 8 s. For stimulus blocks with videos longer than 8 s, the video stimulus was presented once and followed by ~2-s rest period so that the total durations of stimulus blocks can be divided by 2 s (TR). All stimulus blocks were followed by an additional 2-s rest period. Additional 32- and 6-s rest periods were added to the beginning and end of each run respectively. Consequently, the durations of individual runs ranged from 7 min 10 s to 9 min 54 s, and the total duration of all scanning sessions was about 8 hours.

To maximize subject's emotional responses to video stimuli, subjects were allowed to view video stimuli without fixation to let subjects freely focus on any details of events in videos. Subjects were requested to maintain steady fixation on the center fixation spot (0.3 × 0.3 degree) during rest periods to maintain their attention on the screen.

**MRI acquisition**

fMRI data were collected using a 3.0-Tesla Siemens MAGNETOM Verio scanner located at the Kokoro Research Center, Kyoto University. An interleaved T2*-weighted gradient-echo echo planar imaging (EPI) scan was performed to acquire functional images covering the entire brain (TR, 2000 ms; TE, 43 ms; flip angle, 80 deg; FOV, 192 × 192 mm; voxel size, 2 × 2 × 2 mm; slice gap, 0 mm; number of slices, 76; multiband factor, 4). T1-weighted (T1w) magnetization-prepared rapid acquisition gradient-echo (MP-RAGE) fine-structural images of the entire head were also acquired (TR, 2250 ms; TE, 3.06 ms; TI, 900 ms; flip angle, 9 deg; FOV, 256 × 256 mm; voxel size, 1.0 × 1.0 × 1.0 mm).

**MRI data preprocessing**

We performed the MRI data preprocessing through the pipeline provided by fMRIPrep (version 1.2.1; Esteban et al., 2018). For functional data of each run, first, a BOLD reference image was generated using a custom methodology of fMRIPrep. A deformation field to correct for susceptibility distortions was estimated based on fMRIPrep's fieldmap-less approach, and the estimated susceptibility distortion was used to calculate an unwarped BOLD reference for a more accurate coregistration with the anatomical reference. Using the estimated BOLD reference, data were motion corrected using mcflirt from FSL (version 5.0.9; Jenkinson et al., 2002) and then slice time corrected using 3dTshift from AFNI (version 16.2.07; Cox, 1996). This was followed by co-registration to the corresponding T1w image using boundary-based registration implemented by bbregister from FreeSurfer (version 6.0.1; Greve and Fischl, 2009). The coregistered BOLD time-series were then resampled onto their original space (2 × 2 × 2 mm voxels) using antsApplyTrainsforms from ANTs (version 2.1.0; Avants et al., 2008) using Lanczos interpolation.

Using the preprocessed BOLD signals, data samples were created by first regressing out nuisance parameters from each voxel amplitude for each run, including a constant baseline, a linear trend, and temporal components proportional to the six motion parameters calculated during the motion correction procedure (three rotations and three translations). The data were temporally shifted by 4 s (2 volumes) to compensate for hemodynamic delays, were despiked to reduce extreme values (beyond ± 3 SD for each run), and were averaged within each stimulus block (a video presentation period and a subsequent 2-s rest period). The data for all video stimuli were then further z-scored for each voxel. These procedures yielded a total of 2196 data samples each corresponding to each video stimulus. Because the presented video stimulus set happened to include identical videos (15 duplicates), we discarded samples that were presented later in the experiment from each of those duplicates, and used remaining 2181 unique samples in the following analyses.

For visualization of analytical results from the whole cortical areas, we visualized results using flattened cortical surfaces reconstructed from anatomical images of individual subjects. Cortical surface meshes of individual subjects were first generated from the T1w anatomical images using recon-all from FreeSurfer (version 6.0.1; Fischl, 2012). Relaxation cuts were made into the surface of each hemisphere to make flattened cortical surfaces for individual subjects. Functional data were aligned, and were projected onto the surface for visualization using Pycortex (Gao et al., 2012).

**Regions of interest (ROI)**

To define regions of interest (ROI) on cortical surfaces, we used two types of brain parcellations: 1) the whole cortical brain parcellation provided by the Human Connectome Project (Glasser et al., 2016), which delineated a total of 360 cortical areas (180 cortical areas per hemisphere; HCP360; cf., Figures 2 and 3), and 2) the network-based cortical parcellation estimated by intrinsic functional connectivity (Yeo et al., 2011), which delineated 17 networks on the cerebral cortex (cf., Figure 5C). To define ROI masks on individual subject's brain, labels corresponding to brain areas or brain networks originally defined on the standard cortical surface (fsaverage) were converted to the cortical surfaces of individual subjects using FreeSurfer (version 6.0.1; Fischl, 2012). The converted labels were then reinterpolated to 2 × 2 × 2 mm voxels using flirt from FSL (version 5.0.9; Jenkinson et al., 2002). The numbers of voxels in individual ROI masks ranged from 92.4 to 2121.6 for the HCP360 ROIs ($n$ = 360, median = 339.4, five subjects averaged) and from 2121.6 to 12969.8 for the Yeo's 17 network ROIs ($n$ = 17, median = 8290.6, five subjects averaged).

In the analysis of individual cortical areas (Figures 5Band S1A), the visual cortex (VC), temporo-parietal junction (TPJ), inferior parietal lobule (IPL), precuneus (PC), superior temporal sulcus (STS), temporal cortex (TE), medial temporal cortex (MTC), insula, dorsolateral prefrontal cortex (DLPFC), dorsomedial prefrontal cortex (DMPFC), ventromedial prefrontal cortex (VMPFC), anterior cingular cortex (ACC), and orbitofrontal cortex (OFC) were defined based on the following sets of the HCP360 ROI labels on both left and right hemispheres: V1, V2, V3, V3A, V3B, V3CD, V4, V4t, V6, V6A, V7, V8, FST, IPS1, FFC, LO1, LO2, LO3, PH, PIT, MT, MST, VMV1, VMV2, VMV3, and VVC for VC; TPOJ1, TPOJ2, and TPOJ3 for TPJ; PFm, PGi, and PGs for IPL; 7m, v23ab, d23ab, 31pv, 31pd, 31a, and PCV for PC; STSda, STSdp, STSva, and STSvp for STS; TE1a, TE1m, TE1p, and TE2a for TE; EC, and H for MTC; AAIC, MI, PoI1, PoI2, FOP2, FOP3, Ig, and OP2-3 for insula; 9-46d, 46, a9-46v, and p9-46v for DLPFC; d32, 9m, and 10d for DMPFC; 10r, 10v, p32, and s32 for VMPFC; a24, a24pr, p24, a32pr, and p32pr for ACC; and OFC, and pOFC for OFC. Boundaries of these individual brain areas were drawn on the figures of cortical surfaces (e.g., Figures 2C and 4A). For visualization of subareas in VC, lines delineating V1, V2, V3, V4, and others were also drawn on the cortical surfaces. The numbers of voxels in individual cortical brain areas were 19721.2, 2021.2, 5031.4, 2557.0, 2483.8, 5028.8, 1386.4, 2423.8, 4141.6, 3177.4, 1876.4, 2773.0, and 2200.2 for the VC, TPJ, IPL, PC, STS, TE, MTC, insula, DLPFC, DMPFC, VMPFC, ACC, and OFC, respectively (five subjects averaged).

To define ROIs based on the levels of the principal gradient axes, we utilized the principal gradient maps provided from Margulies et al. (2016), which are also originally defined on the standard cortical surface (fsaverage). We first converted original principal gradient maps of the first and second gradient axes to the cortical surfaces of individual subjects using FreeSurfer (version 6.0.1; Fischl, 2012), and then reinterpolated to 2 × 2 × 2 mm voxels using flirt from FSL (version 5.0.9; Jenkinson et al., 2002). The resultant gradient maps registered to individual subjects' brains are shown in Figure 5F (Subject 1), and principal gradient values of individual voxels were used to generate results in Figure 5G and Figure S4E. For each subject, values of the first gradient axis assigned to individual voxels were also used to construct ROI masks that correspond to ten levels (bins) of the first axis, such that a roughly equal number of voxels were assigned to each level (cf., Figure 5H).

To define ROI masks for subcortical areas, including the thalamus, hippocampus, hypothalamus, pallidum, brainstem, caudate, putamen, nucleus accumbens (Brodmann area 34), amygdala, and cerebellum, we used anatomical masks defined by the AAL and the Talairach Daemon provided through the WFU PickAtlas (Maldjian et al., 2003; Lancaster et al, 1997; Lancaster et al., 2000). The anatomical masks, which were originally defined in the stereotaxic space, were transformed to the individual T1w anatomical images, using FreeSurfer (version 6.0.1; Fischl, 2012), and then reinterpolated to 2 × 2 × 2 mm voxels using flirt from FSL (version 5.0.9; Jenkinson et al., 2002). The numbers of voxels in individual subcortical areas were 2738.4, 2763.8, 96.0, 890.4, 5192.8, 2072.4, 2426.4, 596.6, 763.2, and 18712.0 for the thalamus, hippocampus, hypothalamus, pallidum, brainstem, caudate, putamen, nucleus accumbens, amygdala, and cerebellum, respectively (five subjects averaged).

**Video stimulus labeling**

Video stimuli were labeled by multiple types of scores, or features, including two types of emotion scores (34 emotion categories and 14 affective dimensions), 1000 visual object features, and 73 semantic features. Values of these labels were z-scored for each emotion/feature to remove baseline differences across emotions/features (unless otherwise stated).

***Emotion scores***. We used the human emotion ratings of 34 emotion categories and 14 affective dimensions, which were provided from the previous study (see Cowen and Keltner, 2017 for details). The ratings were collected using online experiments via Amazon Mechanical Turk (AMT). For the 34 emotion categories, subjects of the online experiments rated each video in terms of the degree to which it made them feel the 34 emotion categories (100-point scale). The reported scale was converted to 1, when raters scored higher than 0, such that a score for a video from an individual rater become a dichotomous yes/no response. For the 14 affective dimensions, another group of subjects rated each video in terms of its placement along 14 scales of affective dimensions (9-scale Likert scale). Each video was evaluated by multiple raters (9–17 raters), and the ratings obtained for individual emotions from multiple raters were averaged for each emotion to set one mean score for one video for one emotion.

***Visual object features***. For constructing visual object features, we used the Caffe implementation (Jia et al., 2014) of the VGG19 deep neural network (DNN) model (Simonyan and Zisserman, 2014), which was pre-trained to classify 1000 object categories (the pre-trained model is available from https://github.com/BVLC/caffe/wiki/Model-Zoo). The VGG19 model consisted of a total of sixteen convolutional layers and three fully connected layers. To compute outputs by the VGG19 model, all frames of videos were resized to 224 × 224 pixels and provided to the model. The outputs from the last fully connected layer (fc8, 1000 units, before softmax operation) were averaged across all frames within each video to construct a feature vector for a video.

***Semantic features***. We have also collected semantic ratings for the video stimuli according to 73 semantic contents associated with relatively concrete concepts in the video stimuli. The semantic features include objects, scenes, actions, and events (see below for the full list). The data collection was also conducted through online experiments via AMT with 12 raters for each video. Participants rated each video according to whether the video contains each semantic concept by dichotomous yes/no responses. The ratings from 12 raters were averaged to construct feature for individual concept for each video. The full list of the 73 semantic features is as follows: above water scenes, aquatic animals, art, automobiles, babies, birds, black people, blood, boats, bottles/cans, boys, buildings, cartoons, cats, celebrities, cities, clouds, couples, crowds of people, daytime scenes, dead bodies, dogs, elderly people, explosions, fast-moving objects, feces/urine/vomit, fire, flags, food, furniture, genitalia, girls, guns, gymnasiums, hands/feet, historical footage, hospitals, indoor scenes, injuries, insects, land animals, large animals, machines, men, mountains, naked people, nature, nighttime scenes, outdoor scenes, paper, paranormal creatures, people, planes, plants, politicians, reporters, roads, sexual activity, sharp objects, small animals, smoke, snow, soldiers, sports, stores, stunts, television, underwater scenes, vast landscapes, video games, weapons, white people, and women.

**Regularized linear regression analysis**

We used the L2 regularized linear regression (ridge regression) to predict stimulus labels from voxel activity patterns (decoding models) and to predict voxel activity from stimulus labels (encoding models). In the decoding analysis, voxels showing highest correlation coefficients with the target labels in the training data were provided to decoding models constructed for individual labels (with a maximum of 500 voxels). All models were evaluated using 6-fold cross-validation procedure (61 runs data were grouped into five sets of 10 runs and one set of 11 runs data). In each fold of the cross-validation, models for individual labels (decoding) and voxels (encoding) were trained with five sets of data, and the estimated models were used to predict stimulus labels (decoding) and voxel activities (encoding) for left out test set. This was repeated by rotating training-test assignments for 6 times to produce model predictions for all data samples. Then, model performance was evaluated by calculating Pearson correlation coefficients between true and predicted scores of individual labels in the decoding analysis and between measured and predicted brain activities of individual voxels in the encoding analysis. While results shown in this study are based on the 6-fold cross-validation procedure, we have confirmed that differences of the number of fold had little effect on the results.

Ridge regression uses a regularization parameter to constrain the magnitude of the weight coefficients. In decoding analysis, we individually estimated a regularization coefficient for each combination of stimulus labels, ROIs, and subjects (e.g., a single value for “fear” score predictions from V1 activities of Subject 1). In encoding analysis from each label set (category, dimension, visual object, and semantic), we estimated a single value of the regularization coefficient for all voxels in each subject. The regularization parameters were optimized based on the model performances obtained by 5-fold cross-validation procedure (inner loop) nested within training data for the outer loops of the 6-fold cross-validation. In each fold of the nested (or inner) cross-validation loop, models for individual labels (decoding) and voxels (encoding) were trained with four sets of the data (within training data) using each of 20 possible regularization coefficients (log spaced between 10 and 10000), and the estimated models were used to predict stimulus labels (decoding) and voxel activities (encoding) for left out test set (one out of five sets). This was repeated by rotating training-test assignments for 5 times to produce model predictions for all data samples within each fold of outer loops of the cross-validation. Then, model performance was evaluated by calculating Pearson correlation coefficients between true and predicted labels of individual labels in the decoding analysis and between measured and predicted brain activities of individual voxels in the encoding analysis. These procedures were also repeated for each outer loop of the cross-validation to estimate model performances for each fold of outer loops. Then, the regularization parameters producing the maximal model performances in the nested (inner) cross-validation were used for predictions of each fold of the outer cross-validation loops.

**Construction of ensemble decoders**

To construct a decoder that aggregates information represented in multiple brain regions ($n$ = 370), we constructed an ensemble decoder for each emotion by averaging prediction values from multiple decoders (region-wise decoder), each of which was trained with brain activity patterns in each brain region. For each individual emotion, the brain regions used for the aggregations were selected based on the model performances evaluated in a nested cross-validated manner. For each subject and emotion, predictions from region-wise decoders that showed higher decoding accuracy than a threshold ($r > 0.095$, permutation test, $p < 0.01$, Bonferroni correction by the number of brain regions [$n$ = 370]; see Transparent Methods: “Permutation tests” for details) were averaged to construct ensemble predictions. When no brain regions produced decoding accuracy higher than the threshold, the decoder showing the best accuracy was used as a substitute for the ensemble decoder (this was only the case for the “guilt” decoder of Subject 5).

**Video identification analysis**

Identification of emotional experience induced by individual video stimuli was performed via predictions from the decoding (cf., Figures 3A, B and C) and encoding (cf., Figure 4F) analyses. In the decoding and encoding analyses, scores of individual emotions or signal intensity of individual voxels were predicted from observed brain activity patterns or stimulus labels, respectively. Those predictions for individual emotions or voxels were concatenated to construct emotion score patterns or voxel activity patterns. The procedure yielded a total of 2181 predicted patterns corresponding to the presented video clips. Identification was performed in the pairwise manner, in which the video clip was identified between true and false candidates, using the predicted voxel activity pattern or emotion score pattern. The predicted pattern was compared with two candidate patterns: one for the true video and the other for a false video selected from the rest of 2180 videos. The video with a higher correlation coefficient was selected as the identified video. The analysis was repeated for all combinations of the 2181 videos. The accuracy for each video was evaluated by the ratio of correct identification.

**Emotion identification analysis**

To evaluate the inter-subject consistency of brain regions representing individual emotions, identification of emotions was performed via patterns of decoding accuracies obtained from multiple brain regions of different subjects (cf., Figure 2G). In the decoding analysis, we have performed decoding analysis of scores of the 34 emotion categories and 14 affective dimensions, and evaluated decoding accuracies of individual emotions from multiple brain regions (*n* = 370, including both cortical and subcortical regions). The decoding accuracies from multiple brain regions were concatenated to construct a pattern of decoding accuracies (number of elements = 370) as an emotion representation of one subject. For each pair of subjects, accuracy patterns from one subject (test subject) were compared with accuracy patterns from another subjects (reference subject) using all combinations of 34 emotion categories or 14 affective dimensions. For each accuracy pattern of the test subject, the emotion whose accuracy pattern of the reference subject was most correlated with the accuracy pattern of the test subject was selected. This procedure was conducted for all combinations of five subjects (*n* = 20). The identification accuracy was evaluated with various candidate set sizes.

**Slope estimates for performance comparisons**

Comparisons of encoding accuracies from two models (e.g., the emotion category model and affective dimension model; Figures 4B and D) were performed based on the slope angles of best linear fit between two sets of model prediction accuracies of individual voxels. The best linear fit was estimated by Deming regression (Cornbleet and Gochman, 1979; or two-dimensional case of the total least square regression) to accounts for observation errors on both x- and y-axis (e.g., on the category model accuracy and dimension model accuracy; Figure 4B). The slope estimates were converted to angles by first calculating the arctangent of the slopes to obtain angles in radians, and then converting it to degrees. The calculated angles were further subtracted from 45 (degree) to obtain deviations from the parity (Figures 4D and S5D). Statistical significance of the slope estimates was computed based on standard errors of estimated slopes calculated by the jackknife method (two-tailed t-test). For the results of the pooled condition in Figures 4D and S5D, voxels within each brain region were aggregated from all five subjects to calculate slope estimates.

**Dimensionality reduction analysis**

We used Uniform Manifold Approximation and Projection (UMAP) algorithm (McInnes et al., 2018) to perform dimensionality reduction analysis. We applied the UMAP algorithm on emotion category scores (Figure 3D) and brain activity patterns (Figures 6A and B) to reduce the dimensionality of original data into two dimensions.

In the analysis with emotion category scores, we trained a mapping function from original 34 emotion category scores of 2181 video stimuli using correlation distance ($1 - r$) as the distance metric. The trained mapping function was used to project 34-dimensional category scores to two dimensions for both true (Figure 3D left) and decoded (Figure 3D right) emotion category scores.

In the analysis with brain activity patterns, we trained mapping functions based on correlation distances among brain activity patterns to 2181 video stimuli estimated from individual subjects and their average. Before applying UMAP algorithm, voxels associated with emotions were selected based on the results of encoding analysis for individual subjects, in which voxels showing the higher accuracy from the category/dimension emotion models than the visual object and semantic models with significantly high accuracy by the category/dimension emotion models (cf., Figure S4D, including voxels in both cortical and subcortical regions) were selected. The activity patterns of selected voxels were used to construct matrices of correlation distance ($1 - r$) for individual subjects. The estimated distance matrices from individual subjects and their average (2181 × 2181 matrix) were used as inputs to the UMAP algorithm to construct two-dimensional maps of emotional experiences.

In the generated two-dimensional maps, each data sample was colored by a weighted interpolation of the unique colors assigned to 27 distinct emotion categories (Figures 3D, and 6A) or three representative affective dimensions, including valence, arousal, and dominance (Figure 6B).

**Clustering analysis**

We used k-means clustering algorithm to perform clustering analysis with brain activity patterns of individual subjects. The analysis was performed with activity patterns of voxels that were selected based on the results of encoding analyses (see Transparent Methods: "Dimensionality reduction analysis" for the selection criteria of emotion related voxels) using correlation distances ($1 - r$) as metric. The number of clusters ($n$ = 27 in Figure 6) was determined based on the findings in the previous study (Cowen and Keltner, 2017; see Figures S6D and E for results with $n$ = 15 and 50).

**Representational similarity analysis**

The representational similarity analysis (Kriegeskorte et al., 2008) was performed to evaluate the similarity of the representational similarity matrices (RSM) constructed from brain activity patterns and score/feature patterns for each set of features. The RSM was calculated by Pearson correlation coefficients between patterns of voxel activities or scores/features corresponding to individual video stimuli (2181 × 2181 matrix). For calculating the representational similarity between two RSMs (e.g., one from brain activity pattern in a single ROI, and the other from emotion category scores), the off-diagonal elements (triangular part of a matrix) of the RSMs were vectorized and a Pearson correlation coefficient was calculated between those vectors from two RSMs. The analysis was performed using brain activity patterns within individual ROIs defined by the HCP360 parcellation (Glasser et al., 2016). The estimated representational similarities for individual brain regions were averaged across five subjects, and were projected on the cortical surface of Subject 1 (Figure S5).

### QUANTIFICATION AND STATISTICAL ANALYSIS

Two-sided paired t-test was used to examine differences in encoding accuracy from two models (Figures 4C and 5H). ANOVA was used to examine interaction effects between encoding performances from the emotion and semantic models and the levels of principal gradients (Figures 5G and H), and to examine interaction effects between frequency and sorted clusters in the clustering analysis (Figure 6D).

#### Permutation tests

Statistical significance of correlation coefficients (e.g., encoding accuracy in Figure 4A) was computed by comparing estimated correlations (or accuracy) to the null distributions of correlations between two independent Gaussian random vectors of the same length (2181 elements, $n$ = 100,000,000,000; Huth et al., 2016). Resulting $p$-values were corrected for multiple comparisons using the Bonferroni method.

The baseline entropy (cf., Figure 6E) was determined from null distributions of entropies ($n$ = 100,000) calculated from random assignments of the same number of samples ($n$ = 109) into clusters (27 clusters for Figure 6E; 15 and 50 clusters for Figures S6D and E). An entropy was calculated from a histogram by randomly assigning samples into bins. This procedure was repeated for 100,000 times to construct null distributions, and determined the baseline entropy ($p$ = 0.01, Bonferroni correction by the number of emotions times the number of subjects).

### DATA AND SOFTWARE AVAILABILITY

The experimental code and data that support the findings of this study are available from our repository (https://github.com/KamitaniLab/EmotionVideoNeuralRepresentation) and open data repositories (OpenNeuro: https://openneuro.org/datasets/ds002425; Mendeley Data: http://dx.doi.org/10.17632/jbk2r73mzh.1).
